# Adipose Tissue is a Critical Regulator of Osteoarthritis

**DOI:** 10.1101/2020.06.04.134601

**Authors:** Kelsey H. Collins, Kristin L. Lenz, Eleanor N. Pollitt, Daniel Ferguson, Irina Hutson, Luke E. Springer, Arin K. Oestreich, Ruhang Tang, Yun-Rak Choi, Gretchen A. Meyer, Steven L. Teitelbaum, Christine T.N. Pham, Charles A. Harris, Farshid Guilak

## Abstract

Osteoarthritis (OA), the leading cause of pain and disability worldwide, disproportionally affects obese individuals. The mechanisms by which adipose tissue leads to the onset and progression of OA are unclear due to the complex interactions between the metabolic, biomechanical, and inflammatory factors that accompany obesity. We used a murine model of lipodystrophy (LD) to examine the direct contribution of adipose tissue to OA. Knee joints of LD mice were protected from spontaneous or post-traumatic OA, on either a chow and high fat diet, despite similar body weight and the presence of systemic inflammation. These findings indicate that adipose tissue itself plays a critical role in the pathophysiology of OA. Susceptibility to post-traumatic OA was reintroduced into LD mice using implantation of adipose tissue derived from wildtype animals or mouse embryonic fibroblasts that undergo spontaneous adipogenesis, implicating paracrine signaling from fat, rather than body weight, as a critical mediator of joint degeneration.

## Introduction

Osteoarthritis (OA) is the leading cause of pain and disability worldwide, and is associated with increased all-cause mortality and cardiovascular disease (*1–3*). OA is strongly associated with obesity, suggesting that either increased biomechanical joint loading or systemic inflammation and metabolic dysfunction related to obesity are responsible for joint degeneration (*1–3*). However, increasing evidence is mounting that changes in biomechanical loading due to increased body mass do not account for the severity of obesity-induced knee OA (*1–10*). This suggests that other factors related to the presence of adipose tissue and adipose tissue-derived cytokines – termed adipokines – play critical roles in this process (*1, 3, 7, 8*). As there are presently no disease modifying OA drugs available, direct evidence linking adipose tissue and joint health could provide important mechanistic insight into the natural history of OA and obesity and therefore guide the development of novel OA therapeutic strategies.

The exact contribution of the adipokine signaling network in OA has been difficult to determine due to the complex interactions among metabolic, biomechanical, and inflammatory factors related to obesity (*11*). To date, the link between increased adipose tissue mass and OA pathogenesis has largely been correlative (*7, 8, 12*) and as such, the direct effect of adipose tissue has been difficult to separate from other factors such as dietary composition or excess body mass in the context of obesity, which is most commonly derived from excessive nutrition (*3, 7, 8*). Leptin, a pro-inflammatory adipokine and satiety hormone, is secreted proportionally to adipose tissue mass and is the most consistently increased mediator reported in the pathogenesis of obesity-induced OA (*1*). While leptin knockout mice are protected from OA (*7, 8*), the mechanistic role of leptin and other adipokines (*13*) also implicated in OA pathogenesis, remain unknown.

To directly investigate the mechanisms by which fat affects OA, we used a transgenic mouse with lipodystrophy (LD) that completely lacks adipose tissue and therefore, adipokine signaling. The LD model system affords the unique opportunity to directly examine the effects of adipose tissue and its secretory factors on musculoskeletal pathology without the confounding effect of diet (*14, 15*). LD mice completely lack adipose tissue depots, demonstrate similar body mass to WT controls on a chow diet, but exhibit metabolic dysfunction similar to mice with obesity (*12, 14–17*). This allows us to eliminate the factor of loading due to body mass on joint damage and directly test the effects of fat and factors secreted by fat on musculoskeletal tissues. Of particular interest, LD mice exhibit many signs and symptoms that are associated with OA and obesity, including sclerotic bone (*11, 18*), metabolic derangement (*4, 6, 8–10, 19, 20*), and muscle weakness (*3*). Despite these signs and symptoms, we hypothesized that LD mice would be protected from OA due to a lack of adipose tissue, and moreover, that implanting adipose tissue back into LD mice would restore susceptibility to OA – demonstrating a direct relationship between adipose tissue and cartilage health.

## Results

### Male and Female LD mice demonstrate metabolic dysfunction and systemic inflammation

On a chow control diet, both male and female LD mice demonstrated similar total body mass and lean mass compared to their same-sex wild-type (WT) littermate controls (Fig 1A,B). Body fat, measured by both dual-energy x-ray absorptiometry (DXA) and Echo MRI, was significantly reduced in LD mice (Fig 1C), while liver mass was significantly increased in both male and female LD mice (Fig 1D) compared to WT mice. LD mice lacked fat pads throughout the body, including the infrapatellar fat pad normally found in the knee of WT mice *Supplementary Fig 2*.

**Figure 1:**
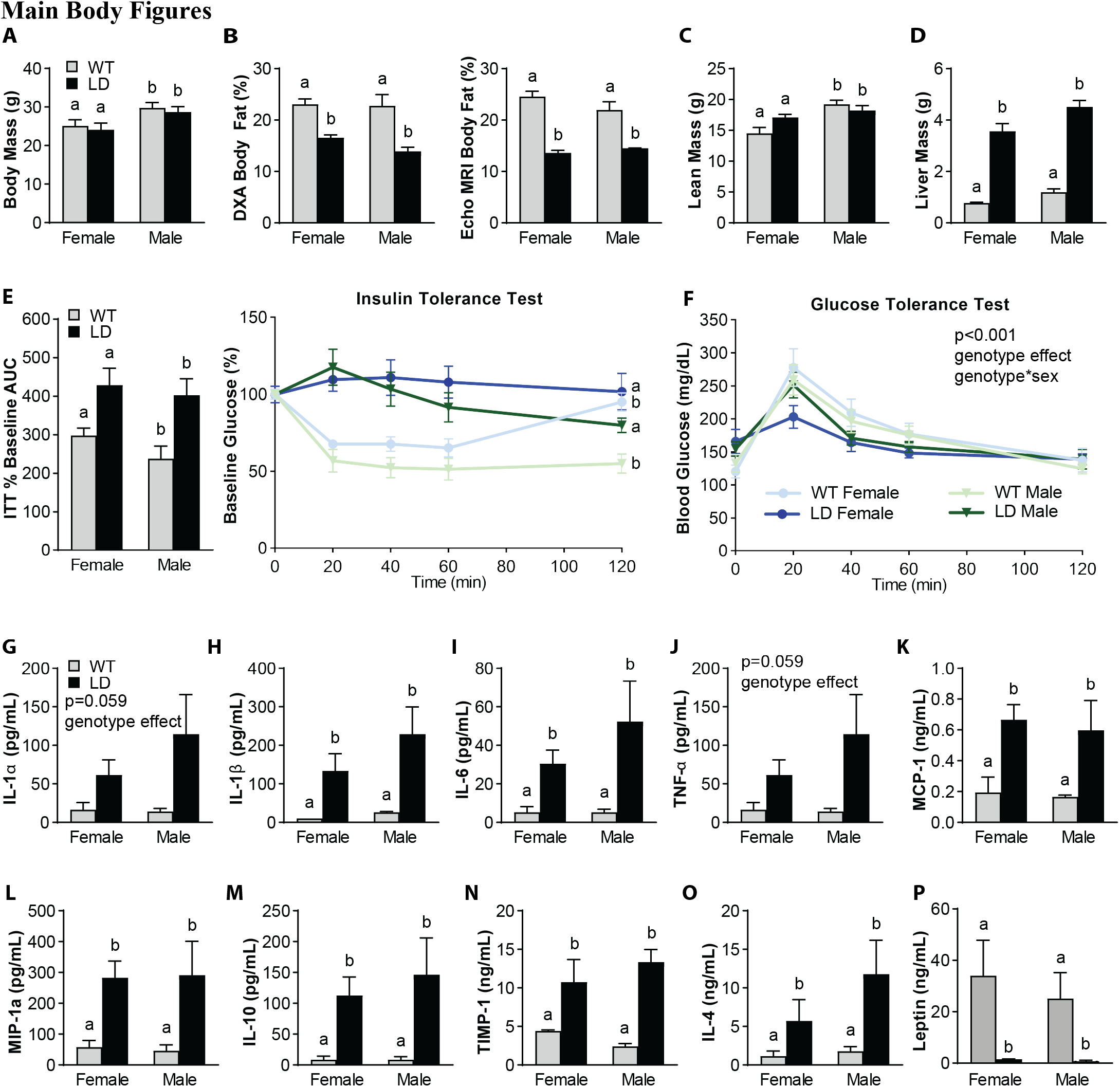
Male and female Lipodystrophic (LD) mice demonstrate metabolic dysfunction and increased pro-inflammatory mediators in serum when compared to wildtype (WT) littermate controls. LD mice have similar body mass (A), but decreased body fat when measured by DXA and Echo MRI (B), and similar lean mass to same-sex WT littermates. LD mice demonstrated increased liver mass (D), insulin resistance and increased area under the curve (AUC) of insulin tolerance tests (E). Glucose tolerance demonstrated a significant main effect of genotype, and genotype*sex. (F). Regardless of sex, LD mice demonstrated increased serum pro-inflammatory (IL-1a, IL-1B, IL-6, TNF-a, MCP-1, MIP-1A) and increased pro-inflammatory mediators (IL-10, TIMP-1, and IL-4) compared to WT littermate controls. LD mice demonstrate near zero levels of leptin in serum (P). N=7-15/group for panels A-F. N=3 for female WT, n=4-7 for other groups for G-O serum outcomes, analysis by 2-way ANOVA with Sidak/Tukey post-hoc; p<0.05 between groups indicated by different letters.

To assess metabolic dysfunction, insulin tolerance tests and glucose tolerance tests were performed. Insulin tolerance tests revealed overt insulin resistance in LD animals compared to WT littermates, as evidenced by a significant increase in blood glucose levels from baseline (Fig 1E). A main effect of genotype and interaction of genotype by sex was significant in glucose tolerance tests, but the area under the curve (AUC) was not different between groups (Fig 1F).

Systemic inflammatory profiles of male and female LD mice indicated similar increases in the pro-inflammatory mediators interleukin-1α (IL-1α), IL-1β, IL-6, tumor necrosis factor alpha (TNF-α), macrophage chemoattractant protein-1 (MCP-1), macrophage inflammatory protein-1 alpha (MIP-1α, Fig 1G-L) compared to WT levels. However, LD mice also demonstrated increased levels of the anti-inflammatory mediators interleukin-10 (IL-10), tissue inhibitor of metalloproteinases-1 (TIMP-1), and interleukin-4 (IL-4) when compared to WT mice (Fig 1M-O). Negligible levels of leptin were detected in serum from male and female LD mice (Fig 1P).

### Chow-fed LD mice demonstrate reduced activity and muscle weakness

To phenotype the activity and metabolic characteristics of the LD mice, we used indirect calorimetry over the period of 36 hours (two dark cycles). No sex differences were observed within each genotype in any of the outcomes assessed. LD mice demonstrate reduced oxygen consumption and energy expenditure (Fig 2A,B). Consistent with a shift toward fat metabolism, LD mice demonstrated a decreased respiratory exchange ratio (Fig 2C) and reduced CO2 production when compared with WT control animals (Fig 2D). LD and WT mice consumed similar amounts of food and water (Fig 2E,F), and exhibited similar energy balance (Fig 2G). LD mice, however, demonstrated decreased locomotor activity (Fig 2H) and ambulatory activity (Fig 2I) compared to WT mice over the observation period. Consistent with this reduced activity, LD mice were significantly weaker by forelimb grip strength assay at 16- and 28-weeks of age (Fig 2J). There was a significant effect of sex and genotype, such that female mice of both genotypes were weaker than their male counterparts.

**Figure 2:**
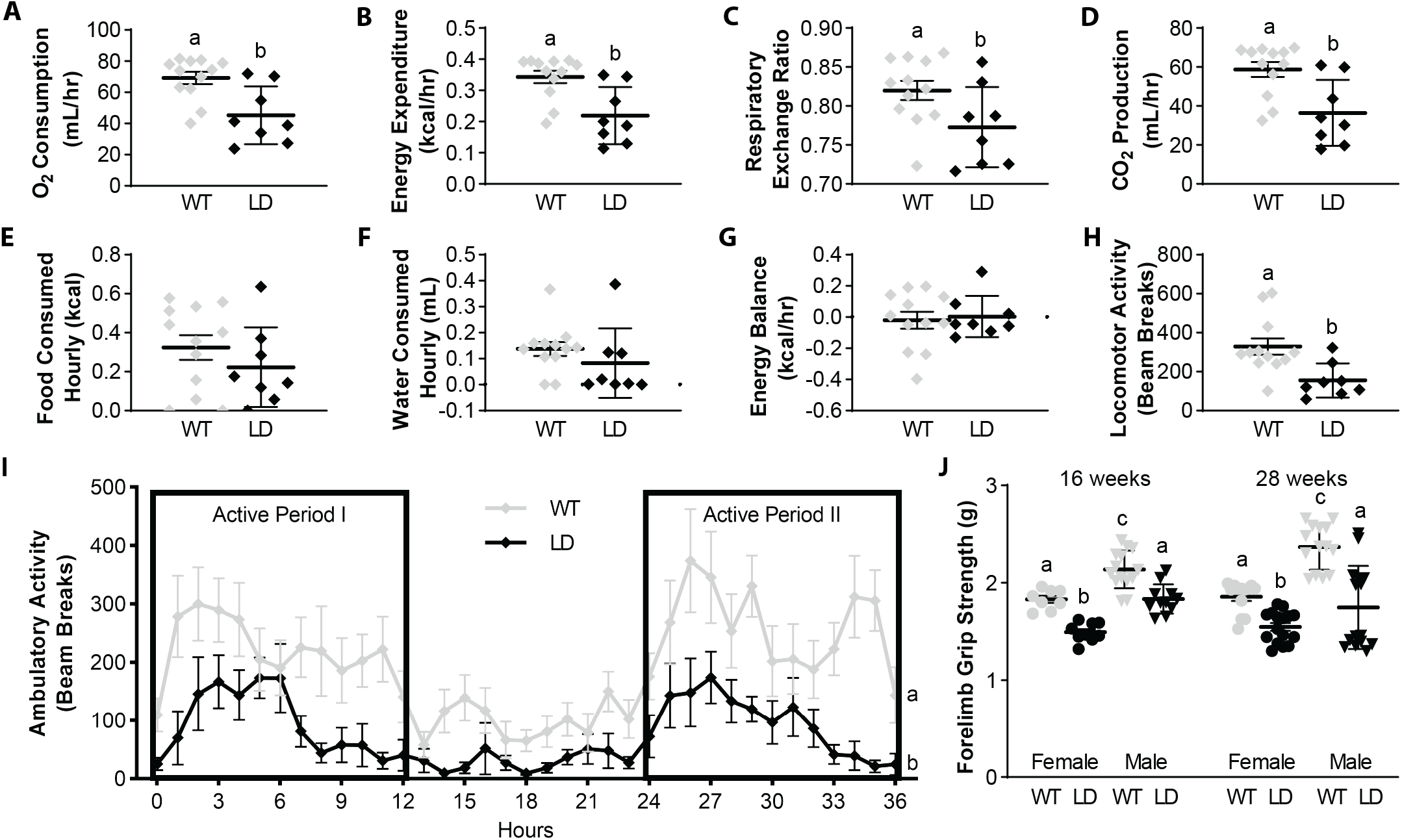
Lipodystrophic mice fed a chow diet demonstrate reduced activity, and muscle weakness despite consuming similar amounts of food and water as WT controls. Using indirect calorimetry, LD mice demonstrate reduced oxygen consumption (A), energy expenditure (B), respiratory exchange ratio (C), C02 production (D), but similar hourly food (E) water (F) and overall average energy balance per hour. LD mice demonstrated lower locomotory (H) and ambulatory (I) activity. No differences were observed by sex, so data were pooled by genotype for comparison. A sex-specific deficit in forelimb grip strength at 16- and 28-weeks, operationalizing muscle strength (J) was observed in LD mice compared to WT Controls. N=7-16/group, A-I general linear modelling using CalR app, 3-way ANOVA with Dunnet’s post-hoc (J), different letters p<0.05.

### LD mice are protected from knee joint damage and pain

Since LD mice show many of the characteristics associated with knee OA, including subchondral bone sclerosis, metabolic disturbance, inflammation, muscle weakness, and decreased activity levels, we evaluated the extent of OA damage in LD knee joints at 28-weeks of age. Unexpectedly, LD mice exhibited significantly less spontaneous knee joint damage in both male and female animals, as measured by Modified Mankin Scores (Fig 3A,D). Next, we challenged the LD animals with a traumatic knee injury using destabilization of the medial meniscus (DMM) surgery. Both WT males and females demonstrated significantly increased damage in DMM joints compared to their non-surgical contralateral limbs, although female WT mice demonstrated less severe OA as compared to male WT counterparts. In contrast, both male and female LD mice were protected from DMM-induced knee joint damage (Fig 3B,E). We observed increased synovitis as defined by histology scoring in DMM knee joints of LD mice compared to contralateral non-surgical knees as well as surgical and non-surgical knee joints of WT mice (Fig 3C,F). There was a trend toward increased osteophytes in DMM limbs of both groups, but the main effect of surgery was not significant (p=0.07, Fig 3B,G).

**Figure 3:**
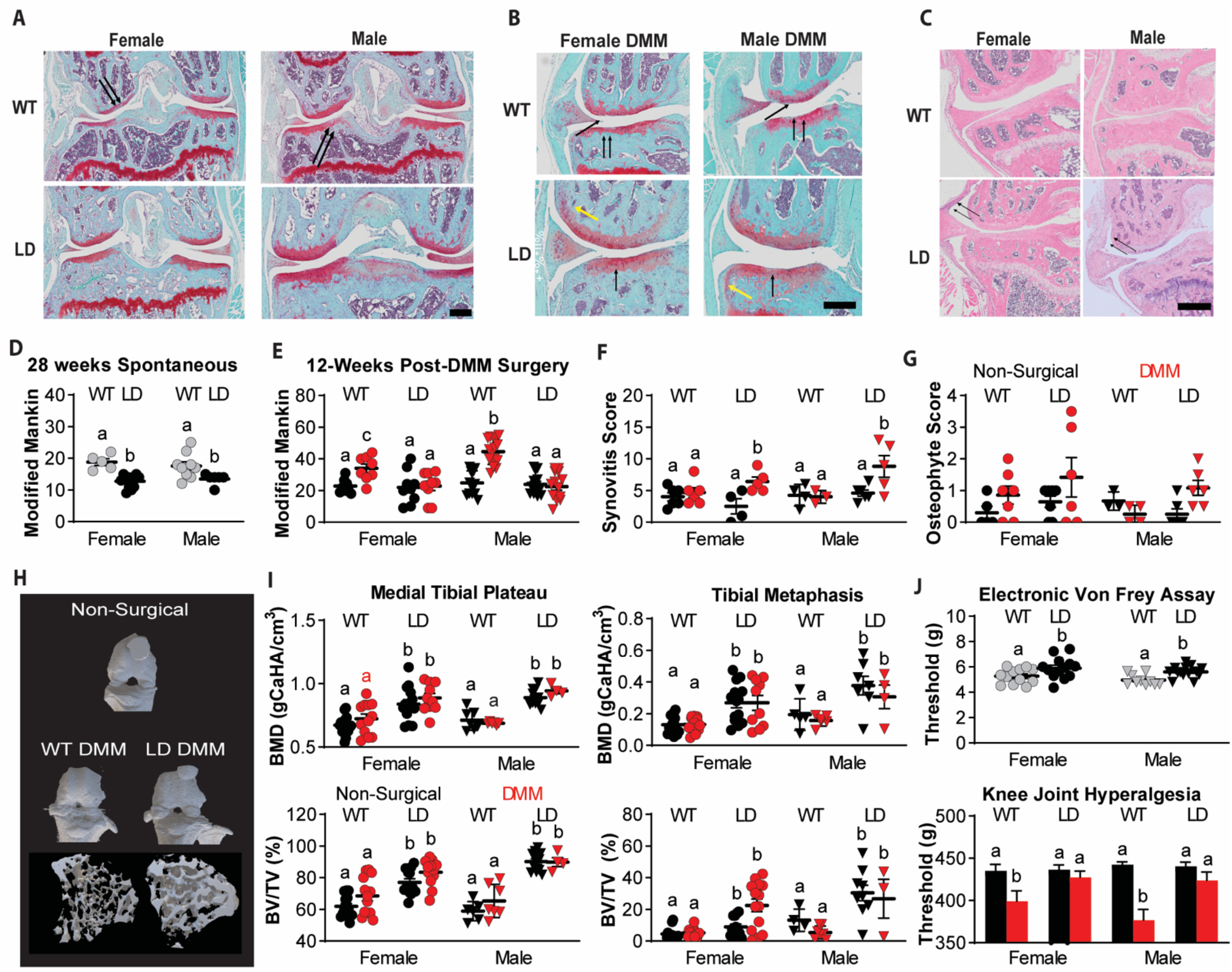
Lipodystrophic mice are protected from spontaneous and post-traumatic osteoarthritis. At 28-weeks, both male and female LD mice demonstrated improvements in Modified Mankin Score (A, D, black arrows indicate cartilage lesion), and when challenged with destabilization of the medial meniscus (DMM) to induce post-traumatic OA, demonstrated similar scores to their contralateral non-surgical limbs (B, E, black indicates non-surgical contralateral limb, red indicates DMM limb). LD male and female mice did demonstrate significant increases in synovitis post DMM (C,F), but similar osteophyte scores compared to WT controls (yellow arrows in B, G). 3D bone renderings of surface and morphology (H). LD genotype drove increases in tibial plateau BMD and BV/TV, as well BMD/BVTV in tibial metaphases (I). LD mice demonstrated protections from the onset of paw allodynia on the DMM limb measured by Electronic Von Frey paw withdrawal assay (J), and hyperalgesia at the knee measured by SMALGO. Data were analyzed as 2-or 3-way ANOVA with either Sidak/Tukey/Dunnet post-hoc; different letters p<0.05.

As sclerotic subchondral bone is widely implicated in OA onset and pathogenesis, we assessed bone microstructure of LD mice and compared it to WT mice using microCT analysis. LD genotype, but not sex or surgery, drove significant increases in male and female bone mineral density in the medial tibial plateau (Fig 3H,I). In addition to genotype, a significant main effect of surgical intervention was observed in medial tibial plateau BV/TV, and a significant interaction between sex and genotype was seen. Tibial metaphysis BMD also demonstrated significant main effects of sex and genotype. Tibial metaphysis BV/TV demonstrated a significant main effect of sex and genotype, and significant interactions between sex*limb, and sex*genotype were observed. Trabecular thickness, trabecular number, BMD and BV/TV for the lateral tibial plateau, medial femoral condyle and the medial tibial plateau generally demonstrated significant effects of genotype. Detailed site-specific outcomes for each bone microstructure parameter are described and compared in *Supplementary Table 2*. Generally, LD mice demonstrate a strong increased bone mass phenotype when compared to WT littermate controls.

We performed tests to quantify knee hyperalgesia and tactile allodynia to evaluate if LD mice develop increased pain sensitivity as a consequence of DMM. LD mice exhibited significantly higher paw-withdrawal thresholds when the DMM paw was subjected to an Electronic Von Frey assay to determine tactile allodynia (Fig 3J) compared to WT animals. Furthermore, LD mice maintained similar pressure-pain thresholds in both the surgical and non-surgical contralateral knee while WT DMM knee joints demonstrated knee hyperalgesia.

### High fat diet does not exacerbate the pro-inflammatory profile of LD mice

Given the inability of LD mice to store fat, we next sought to determine if a high fat diet (HFD) would alter the metabolic disturbance and inflammation in LD mice. Beginning at weaning, LD and WT male and female animals were fed either a 60% kcal fat diet or 10% kcal fat chow diet. In WT mice, body mass and body fat were increased with HFD. However, LD mice fed HFD had no change in body mass or body fat, and were similar to LD and WT mice on a chow diet (Fig 4A,B). Lean mass at 28-weeks was significantly reduced in both the LD and WT HFD-fed animals (Fig 4C) as compared to both genotypes on the chow diet. Liver mass was elevated in LD mice compared to WT mice, but not significantly increased between chow and HFD (Fig 4D).

**Figure 4:**
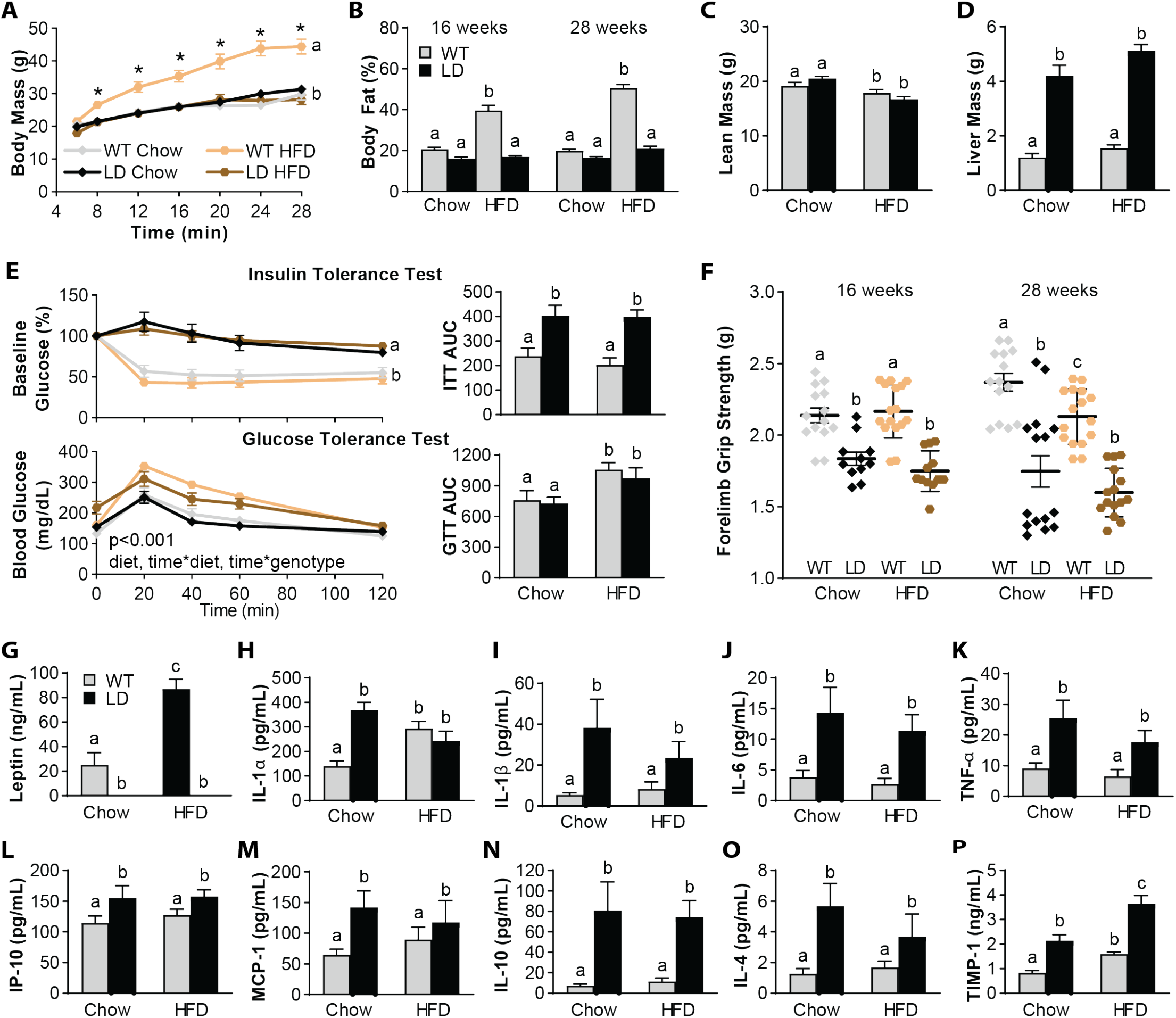
Superimposition of a high-fat diet (HFD) on fat-free lipodystrophic mice does not increase the metabolic derangement of lipodystrophic mice. Body mass (A) and body fat (B) over time was only increased in WT animals fed HFD, but not LD fed HFD. Lean mass was decreased in both groups fed HFD (C), but liver mass (D) was increased in LD mice to a similar extent even on HFD, when compared to WT. Insulin resistance by insulin tolerance test was not affected by HFD (E), and differences were only observed by genotype, as in the area under the curve (AUC). Glucose sensitivity was reduced by HFD, as indicated in the AUC. Muscle weakness was also decreased in LD mice, and at 28-weeks, decreased in HFD-fed WT animals when compared to their 16-week timepoint (F). Serum leptin was increased in WT animals fed HFD, but still completely absent in LD mice regardless of diet (G). Levels of pro-inflammatory mediators that were elevated in LD mice were not further elevated by HFD (IL-1a, IL-1B, IL-6, TNF-a, IP-10, MCP-1, H-M). Levels of anti-inflammatory mediators IL-10 and IL-4 were similarly not increased by HFD in LD mice, but TIMP-1 was significantly increased (N-P). Data were analyzed as 2-or 3-way ANOVA with either Sidak/Tukey/Dunnet post-hoc; different letters p<0.05, n=10-18/group.

In LD mice, a HFD did not worsen insulin tolerance in either genotype, such that HFD LD and WT mice were similar to their chow WT counterparts (Fig 4E,F). Glucose tolerance, however, was significantly reduced in both LD and WT groups, as demonstrated both in the time-course and AUC curves (Fig 4E). HFD did not further decrease muscle weakness measured by forelimb grip strength in LD mice, but did reduce grip strength in WT HFD-fed mice (Fig 4F). Circulating leptin was increased in HFD-fed WT animals compared to chow-fed WT, but leptin remained undetectable in the serum of HFD-fed LD mice. (Figure 4G). HFD did not result in increased circulating pro-inflammatory mediators in LD mice, as levels for IL-1α, IL-1β, IL-6, TNF-α, IP-10, and MCP-1 were similar to chow-fed LD levels (Fig 4H-M). IL-10 and IL-4 remained elevated in LD animals with HFD, whereas TIMP-1 was further elevated in HFD-fed LD mice, such that it was increased compared to both chow-fed LD mice and both WT groups (Fig 4N-P). Further differences in serum profiles are detailed in *Supplementary Table 1*.

### High-fat fed LD mice are protected from OA

We then aimed to determine if high fat feeding could override the protection against joint damage observed in chow-fed LD mice. Both male and female HFD-fed LD mice remained protected from OA as assessed by Modified Mankin Scores (Fig 5A,B). Both HFD-fed groups demonstrated significantly increased osteophyte scores in the DMM limbs (Fig 5C). Synovitis scores, however, were significantly increased in chow-fed LD DMM limbs, and there was a trend toward a decrease in synovitis of HFD LD DMM limbs (Fig 5D). Significant negative associations were observed between serum levels of IL-10 (r^2^=0.42, p<0.001), TIMP-1 (r^2^=0.38, p<0.001), IL-4 (r^2^=0.16, p<0.035) and Modified Mankin Scores of histological knee joint damage when Pearson correlations were calculated. No significant relations were observed between pro-inflammatory mediators like IL-6 (r^2^=0.06, p=0.22) and Modified Mankin Scores.

**Figure 5:**
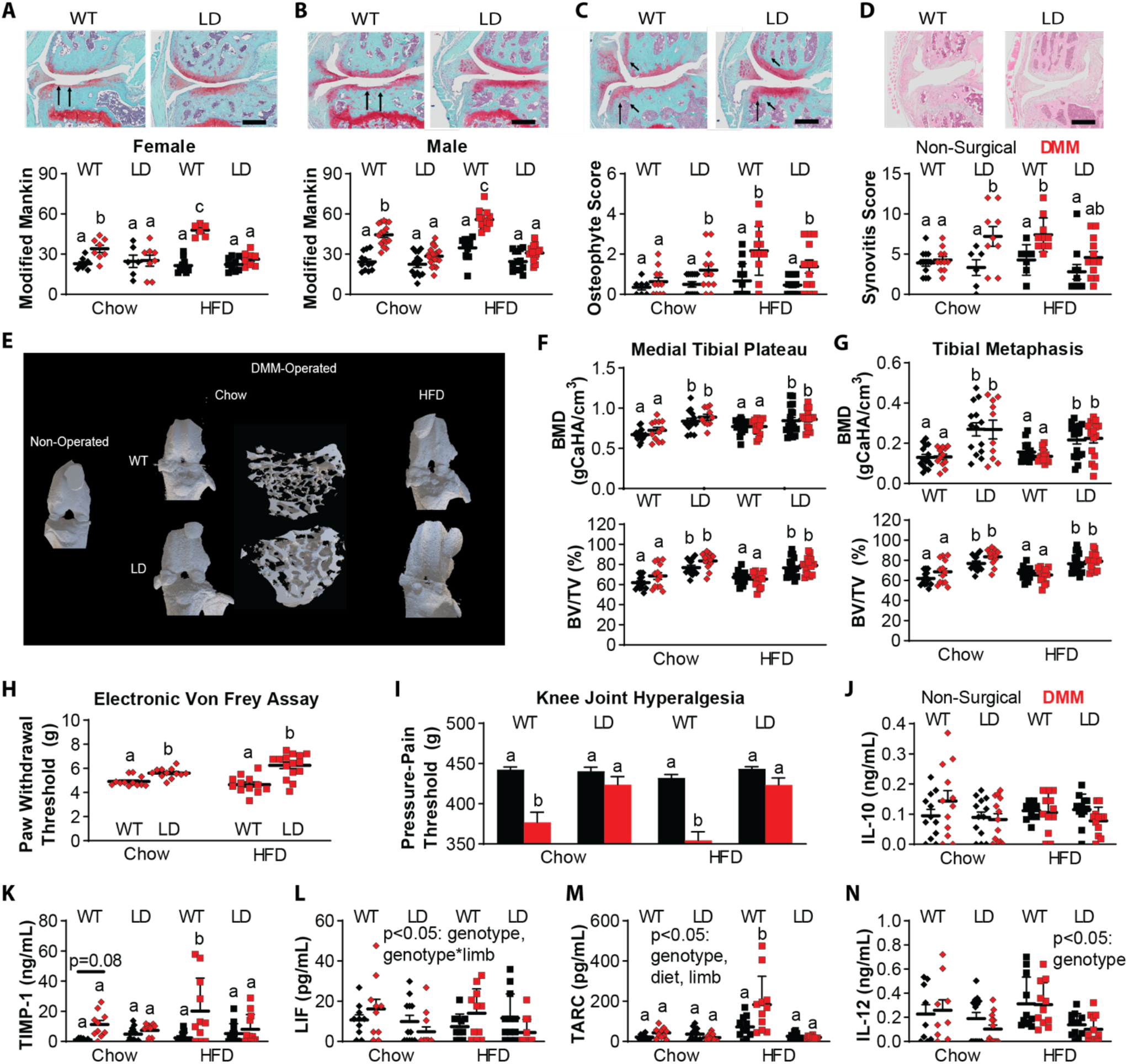
High fat feeding does not override protection from post-traumatic OA in LD mice. WT HFD-fed male and female animals demonstrated increased Modified Mankin Scores when compared to both chow-WT DMM limbs and contralateral non-surgical controls (A-B, black indicates non-surgical contralateral limb, red indicates DMM limb). Regardless of diet or DMM surgery, LD mice demonstrated similar Modified Mankin scores to their non-surgical contralateral limbs. HFD-fed LD mice did demonstrate a trend toward a reduction in synovitis in the DMM limb (C). Both HFD WT and LD mice demonstrated increased osteophyte scores in the DMM limb (D, black arrows). 3D bone images from knee joint microCT and trabeculae from tibial metaphysis (E). Genotype drove the differences in bone mineral density and bone volume/total volume of medial tibial plateau and tibial metaphases (F-G). LD mice on both diets demonstrated protection from the onset of knee joint hyperalgesia measured by SMALGO and paw withdrawal threshold at the paw measured by Electronic Von Frey (H-I) compared to WT controls in DMM limbs. Synovial fluid levels for anti-inflammatory mediators IL-10 and TIMP-1 were similar by joint, genotype, and diet (J-K). Reductions were observed in DMM LD joint synovial fluid levels for LIF, TARC (L-N). Remaining synovial fluid profiles are reported in Supplementary Data. Data were analyzed as 2-or 3-way ANOVA with either Sidak/Tukey/Dunnet post-hoc, Pearson correlation; different letters p<0.05, n=10-18/group.

As with chow-fed animals, genotype was the principle driver of increased BMD and BV/TV in both the medial tibial plateau and the tibial metaphyses (Fig 5E-G), as well as the lateral tibial and femoral compartments (*Supplementary Table 2)*. Medial tibial plateau BV/TV also demonstrated a significant main effect of limb, and an interaction between limb*diet. Tibial metaphysis BV/TV demonstrated significant main effects of limb in addition to genotype, and significant interactions between limb*diet, limb*genotype, and limb*diet*genotype. Additional bone microstructure outcomes are detailed in *Supplementary Table 2*.

LD animals fed chow and HFD were protected from hyperalgesia, as measured by the decreases in paw withdrawal thresholds (Fig 5H) and pressure-pain thresholds (Fig 5I) observed in WT animals. Despite consistent increases in IL-10 in serum of LD mice on both diets, we did not observe changes in synovial fluid levels for IL-10 (Fig 5J). In WT animals, HFD-feeding was associated with increased synovial fluid TIMP-1 levels in the limb that underwent DMM (Fig 5K). There were significant decreases in synovial fluid levels of LIF, TARC, and IL-12 (Fig 5L-N) in DMM limbs of LD mice. Synovial fluid profiles are detailed in *Supplementary Table 3*.

### Transplantation of Adipose Tissue Restores Susceptibility to OA in LD Mice

To determine if adipose tissue is a critical regulator of OA, we transplanted adipose tissue into chow-fed LD mice using one of two methods. First, subcutaneous and visceral adipose tissue from WT mice was implanted subcutaneously on the dorsum of LD mice (termed “WT fat rescue”). A separate cohort received cell-based injections of primary mouse embryonic fibroblasts (MEFs) harvested from WT pregnant dams (termed “MEF rescue”). MEF injections, delivered subcutaneously to the sternal aspect of mice, later form a fat depot approximately 21 days post injection as previously described in this mouse model (*16, 17*).

Both groups of LD fat transplant animals were challenged with DMM surgery, and histological assessment revealed that both WT fat rescue and MEF rescue animals demonstrated significant increases in DMM joint damage when compared to non-surgical contralateral limbs. The severity of joint damage in either of the rescued LD mice was similar to DMM damage in WT control animals (Fig 6A), suggesting that fat implantation reintroduces susceptibility to OA. Fat implant size was similar between groups (WT fat rescue 0.95±0.1g, MEF rescue 0.80 ±0.07g, p=0.18), and mice receiving a fat implant showed similar body mass in WT fat rescue (24.2±0.7g) and MEF rescue (23.0±0.2g) mice as compared to the average body mass of chow-fed LD (26.3±1.2g, by 1-way ANOVA p=0.12) mice. The size of the implant demonstrated a weak correlation with Modified Mankin Score (r^2^=0.22, p=0.0176). When analyzed separately, the size of WT fat rescue implant size demonstrated no significant relationship with damage, while the MEF rescue implant demonstrated a strong significant relationship (r^2^=0.66, p<0.001). Across all chow-fed animals, with and without transplant, there was no significant correlation between body mass and Modified Mankin Score (r^2^=0.028, p=0.203).

**Figure 6:**
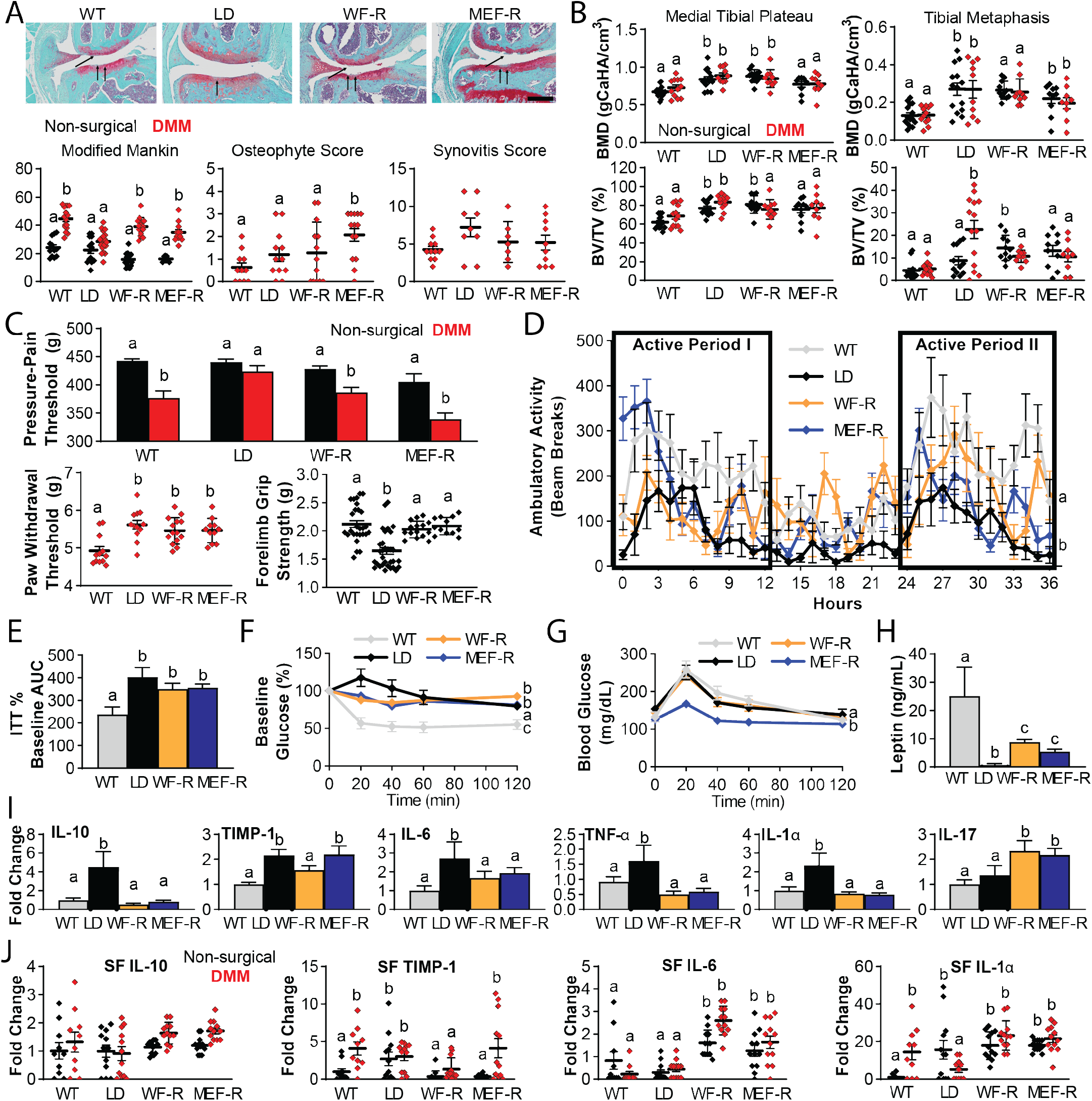
Transplantation of fat restores susceptibility to OA damage and partially corrects the metabolic derangement in LD mice. Both wildtype mature adipose (WF-R) transplant and primary mouse embryonic fibroblast (MEF-R) transplant in LD mice restores increased Modified Mankin Scores in DMM limbs (A, black indicates non-surgical contralateral limb, red indicates DMM limb). Osteophyte scores were significantly increased in MEF-rescue animals compared to the other groups in DMM limbs. Synovitis scores were similar across groups in DMM limbs. Medial Plateau BMD and BVTV in contralateral limb was reduced to WT levels in MEF-rescue animals only (B); WT fat-rescue animals’ DMM limbs demonstrated significant reductions in BV/TV to WT levels. Both transplant types restored susceptibility to knee hyperalgesia (C), but not paw allodynia in the DMM limb. Forelimb grip strength was also restored to WT levels in both transplant groups. Restoration of activity levels was also observed in both transplant groups (D). While both fat transplants demonstrated overall improvements in insulin resistance (E-F), the AUC was not significantly different from the LD mice. WT fat-rescue, however, significantly improved glucose tolerance (G). Leptin ELISA demonstrated that both fat transplants reintroduce circulating leptin levels to LD mice (H). Serum levels for IL-10 were reduced in both transplant groups compared to LD mice (Q), but serum TIMP-1 was only reduced in WT fat-rescue mice (I). Other pro-inflammatory mediators (IL-6, TNF-a, IL-1a, Z-AB), were also improved in transplant mice, whereas IL-17 was significantly elevated. Synovial fluid levels for IL-10 were similar across groups (J), whereas TIMP-1 levels in the DMM limbs were elevated to WT levels in MEF-rescue LD mice. Synovial fluid levels for IL-6 and IL-1a were increased in WT fat-rescue and MEF-rescue animal DMM limbs. Data were analyzed as 1, 2-or 3-way ANOVA with either Sidak/Tukey/Dunnet post-hoc tests, different letters p<0.05, n=7-18/group.

Osteophyte scores were only increased in MEF treated LD animals (Fig 6A), while synovitis scores were similar across WT, LD, and the two transplant groups (Fig 6A). Increases in BMD were reversed to WT levels in the medial tibial plateau of MEF rescue LD animals, while BMD was similar in both limbs from WT fat rescue animals to LD mice (Fig 6B). Medial tibial plateau BV/TV was reversed to WT levels in MEF-rescue LD mice, but only in the DMM limb of WT fat rescue mice (Fig 6B). Tibial metaphysis BMD was partially reversed to WT levels in MEF rescue animals, but similar to LD mice in WT fat-rescue mice (Fig 6B). Tibial metaphysis BV/TV was reduced to WT levels in the DMM limbs of both fat transplanted groups, while the non-surgical contralateral limbs were similar to DMM limbs in LD mice (Fig 6B). Data for lateral tibial plateau, medial femoral condyle, and lateral femoral condyle, as well as trabecular thickness and trabecular number, can be found in *Supplementary Table 2*.

Knee hyperalgesia in response to DMM was restored in WT fat-rescue and MEF-rescue LD animals to similar levels measured in WT DMM limbs (Fig 6C), while DMM limb paw-withdrawal threshold remained similar in both fat transplant groups to the values measured in LD animals (Fig 6C). Forelimb grip strength was also restored in both WT fat rescue and MEF rescue animals (Fig 6C).

Several aspects of the systemic metabolic profiles were improved in both WT fat-rescue animals. Reduced locomotor activity was also reversed in both fat transplant groups, such that the WT fat rescue and MEF rescue groups showed activity profiles that were similar to WT animals and were increased compared to LD animals (Fig 6D). Insulin tolerance curves were significantly reduced in both WT fat rescue and MEF rescue groups, but the AUC of these curves remained similar to the AUC of LD mice (Fig 6E,F). Glucose tolerance curves were significantly improved in WT fat rescue animals, but curves from MEF rescue animals were similar to WT and LD animals (6G). Serum leptin was restored in both MEF rescue and WT fat rescue LD mice, to approximately 20-30% of endogenous circulating levels in WT mice (Fig 6H).

Fat transplantation reduced the circulating levels of anti-inflammatory mediators IL-10 and TIMP-1 in LD mice to WT levels (Fig 6I). Pro-inflammatory mediators IL-6, TNF-α, and IL-1α were also reduced in WT fat rescue and MEF rescue animals, but IL-17 was significantly increased when compared to both LD and WT treated animals (Fig 6I). Serum levels for IL-10 demonstrated a modest, but significant negative relationship with Modified Mankin Score when the fat transplanted animals were included in the correlation analysis (r^2^=0.26, p<0.001). Serum differences due to fat transplantation are listed in *Supplementary Table 4*.

Despite the alterations we observed in the serum profiles of fat transplant animals, we did not observe any differences in IL-10 concentration in the synovial fluid (SF, Fig 6J). SF levels for TIMP-1 in MEF-rescue animals were similar to those observed on contralateral and DMM limbs of WT mice; however, WT fat-rescue levels were decreased in both limbs, such that they were similar to the levels observed in the WT contralateral limb (Fig 6J). Levels for IL-6 were significantly increased in both fat transplant groups compared to LD and WT limbs, whereas SF IL-1α levels were decreased in LD DMM limbs, but similarly increased in the both limbs of WT fat rescue and MEF rescue LD animals (Fig 6J). A list of SF changes in WT fat rescue and MEF rescue animals can be found in *Supplementary Table 5*.

## Discussion

While several studies have illustrated important relationships between adiposity, low-level systemic inflammation, and OA, the direct role of adipose tissue on joint health has been difficult to verify due to the complex interactions among adiposity, body weight, dietary content, and inflammation (*1, 3–5, 7–10, 20–23*). Here, we demonstrate that knee joints from mice lacking adipose tissue are protected from spontaneous or injury-induced OA, even when fed a high-fat diet. These studies demonstrate a profound role for adiposity, rather than the high-fat diet *per se*, in exacerbating the severity of OA injury. Of particular note was the finding that susceptibility to post-traumatic OA can be restored in LD mice after implantation of a small adipose graft or injection of MEFs, which form a functional fat depot. These experiments implicate adipose tissue, and presumably, factors secreted by adipose tissue as a mediator of joint degeneration, rather than body weight, bone quality, muscle strength, metabolic derangement, or diet. Furthermore, these data support the notion that OA can be influenced and controlled by tissues outside of the joint organ system, as the addition of metabolically healthy fat can increase susceptibility to OA.

Notably, while LD mice lack adipose tissue, which is thought to serve as a local source of pro-inflammatory mediators and adipokines within articulating joints (*23, 24*). LD mice exhibited several characteristics traditionally thought to contribute to the onset and progression of cartilage damage associated with obesity. Despite exhibiting low-level systemic inflammation (*4, 7–9*), metabolic disturbance (*1, 3*), insulin resistance (*22*), sclerotic bone (*11*), muscle weakness (*3, 6*), reduced activity (*25*), synovial inflammation (*1, 26*), LD mice were protected from cartilage damage. Because increased pain and hyperalgesia are the primary clinical symptoms of OA (*27*), we also assessed two pain-related outcomes: knee hyperalgesia and tactile allodynia, and observed that consistent with a protection against structural damage, LD mice are also protected from the onset of these pain-related outcomes with DMM. Remarkably, the transplantation of a small amount of adipose tissue mitigated or reversed these contributing factors and restored susceptibility to cartilage damage and knee hyperalgesia in transplanted LD mice. This controlled model of lipodystrophy confirms a direct relationship between adipose tissue and the onset and progression of OA, independent of biomechanical and metabolic contributors, although the specific factors involved in this protection remain to be clarified. Importantly, this model provides a system that will us to disentangle the contributions of specific adipokines to cartilage damage.

Because LD mice demonstrate protection from cartilage damage, it is likely that systemic or local inflammatory mediators may be playing a protective role from knee joint damage and could be employed as therapeutic avenue for OA. Adipose tissue could drive OA susceptibility directly through adipokines, through changes in systemic inflammation that ultimately manifest locally within the joint, or indirectly through another signaling axis absent in LD mice. In this regard, leptin is the most consistently increased mediator reported in obesity-induced OA (*7, 8, 24, 28*) and can drive OA-associated inflammation (*28*). Obesity is associated with increased circulating levels of leptin thought to arise in response to leptin resistance (*3*). Leptin knockout mice and LD mice, which completely lack leptin, are protected from OA (*7, 8*). Consistent with previous studies, we observed that metabolic derangement and adipokine signaling can be rescued in the LD mice by the implantation of a surrogate fat pad derived from a mouse embryonic fibroblasts cell-based injection (MEFs), or through mature adipose tissue implantation (5,15), that restores between 20-30% of normal leptin levels in LD-transplanted mice. However, inhibition of leptin has failed in the clinical setting, and directly exposing cartilage explants to leptin does not induce cartilage damage (*7*). Therefore, we posit that the critical mediator involved in this protection is a downstream target of leptin but ongoing leptin-targeted studies will further elucidate the specific role of this adipokine.

LD mice demonstrate consistently higher levels of pro-inflammatory mediators, that appear to be balanced by high levels of anti-inflammatory mediators (IL-10, TIMP-1, IL-4). Furthermore, our cytokine array data on LD mice identified increased serum levels of IL-10, a potent anti-inflammatory molecule that has been proposed as an OA therapeutic and pain reliever (*29, 30*). In LD mice receiving adipose transplantation, where susceptibility to OA was restored, we observed a decrease in circulating serum levels of IL-10. IL-10 concentrations demonstrated a negative association with cartilage damage, which is consistent with reports of IL-10 as a potential beneficial therapeutic agent for OA (*29, 30*), and could be fundamental to the potential systemic basis of cartilage protection observed in LD mice. Moreover, IL-10 can functionally mimic leptin due to high structural homology (*31*), and reduces insulin resistance and high-fat diet-induced inflammation in models of obesity without off-target pro-inflammatory consequences (*32*). It is also possible that IL-10 is driving the increases in TIMP-1 that were observed in serum and synovial fluid of LD mice, given that IL-10 is known to induce TIMP-1 and reduce the activity of matrix metalloproteinases (*33*). Future work will determine if sustained increases in circulating levels of IL-10 is indeed a key protective mechanism by which LD mice remain resistant to the onset of cartilage damage.

Adipose transplantation increased circulating levels of IL-17 compared to LD and WT mice, which has previously been implicated in OA pathogenesis (*26*). It is possible that the systemic increases in IL-17 from MEF-rescue and WT fat-rescue are derived from a host-response (*34*); however, as the MEF-rescue is delivered as a cell-based therapy that ultimately forms a tissue that integrates well into the host (*16, 17*), it is difficult to determine if that is the explanation for increased IL-17 in these animals. However, IL-17 has been shown to regulate adipogenesis, glucose homeostasis (*35*), and obesity, so the IL-17 increases observed in MEF-rescue and WT fat-rescue serum may be instead related to the integration and lipid storage that results in implanted LD mice. Several studies have demonstrated that IL-17 contributed to OA pathogenesis (*26*). Ultimately IL-17 could be a key factor contributing to the restoration of OA susceptibility in LD mice upon fat transplant, as others have reported IL-17 as a potential biomarker for a distinct subset of inflammatory OA patients with end stage disease (*36*).

It is important to note that LD mice also lack other signaling pathways, such as the alternative complement pathway. LD mice have an absence of adipsin, or complement factor D, which cleaves factor B to initiate alternative complement pathway activity (*12*). Previous studies have demonstrated a protective effect of adipsin deficiency on both spontaneous and post-traumatic OA pathogenesis (*13, 37*), so it is unclear if adipsin deficiency is the protective adipokine mechanism modulating joint damage in the present studies. Our ongoing work is interrogating the role of adipsin knockout animals in the presence of high fat diet and osteoarthritis to determine if the absence of adipsin signaling in particular protects against cartilage damage in the LD mouse.

While we observed several differences by genotype and diet in the synovial fluid of LD mice that are concordant with OA pathogenesis (*26*), we did not observe overt structural damage to LD knee joint cartilage. Specifically, we observe reduced levels for leukemia inhibitory factor (LIF) and interleukin 12 (IL-12) in the DMM limbs of LD mice, and a concordant increase in IL-1α and IL-6 in MEF-rescue and WT fat-rescue LD mice. This implicates a role for the IL-1 and IL-6/IL-12 family of cytokines in protection and susceptibility to OA, which has been postulated as key therapeutic target networks for OA (*26*). In addition, we observed decreased levels for TARC/CCL-17 in LD DMM limbs, which is of interest as CCL-17 blockade has been demonstrated as a therapeutic candidate for OA structure and pain (*38*). In the fat rescue groups, we observed increased circulating levels of TARC/CCL-17, but decreased SF levels for TARC/CCL-17. IL-4 is another promising anti-inflammatory candidate for cartilage protection under active investigation, and here we observed increased IL-4 in LD mice (*39*). However, as IL-4 can also induce TARC/CCL-17 via STAT6, there may be off-target pro-inflammatory consequences of overexpressing IL-4 therapeutically in situations of low-level chronic inflammation (*40*) which may detract from the therapeutic potential of IL-4 in this context. Future studies will evaluate the direct influence of IL-6/IL12 axis and TARC/CCL-17 signaling in the conference of protection from OA in LD mice.

A striking absence of anti-inflammatory signatures were found in the synovial fluid of LD mice when compared to WT mice. There are a few plausible explanations for this finding. First, evaluating data cross-sectionally at the terminal endpoint of 12-weeks post-DMM does not allow for temporal assessment of changes in the systemic or local inflammatory environment and it is possible that critical changes in the synovial fluid that contribute to cartilage protection occur soon after DMM and are not detectable at terminal endpoint. A second possibility is that the systemic environment, which is directly affected by the absence and implantation of adipose tissue, is mitigating and driving OA disease pathogenesis in this LD model system. This proposition is consistent with several studies that demonstrate increased body mass *per se* does not explain obesity-associated knee OA, but that in fact, associations with systemic pro-inflammatory mediators and other tissues (e.g., muscle) provide robust associations with OA pathogenesis (*1, 5, 7, 8, 24, 28*), which is the opposite of what we observed in the LD mice. As such, we postulate that the strong anti-inflammatory signature evidenced in LD mice, which is reversed upon adipose implantation, is a key driver of this protection, and that systemic overexpression of anti-inflammatory mediators may be a useful therapeutic approach for chondroprotection with injury or obesity.

An important finding in these studies is the remarkable presence of bone sclerosis in concordance with healthy cartilage in LD mice, which was subsequently reversed with fat transplantation. Many studies have hypothesized that subchondral sclerosis is a pathological characteristic of OA (*11, 18*), even proposing that subchondral bone remodeling is a primary driver of cartilage damage (*18*). However, our findings clearly demonstrate that these factors are biologically separable, and healthy cartilage can persist even when challenged with trauma, in the presence of marked subchondral sclerosis and increased bone density throughout the joint. Similarly, the reversal of muscle weakness observed in LD mice with fat implantation contradicts many studies that have suggested a key role for muscle weakness in the pathogenesis of OA (*3, 9*). This includes recent work from our laboratory that demonstrates risk for post-traumatic OA can be mitigated by enhancing muscle structure and strength and correcting metabolic derangement using a gene therapy approach (*6*). Future studies will aim to resolve these issues and disentangle these risk factors for obesity and OA using the LD mouse platform and time-course studies. These insights may contribute to our evolving understanding of mechanical and biological tissue cross-talk, both within and between the joint organ system, to help determine what constitutes compensatory versus pathological loading environments. Using this insight, we can begin to unravel intrinsic mechanisms that other tissues of the joint organ system or intrinsic systemic tissues may employ in efforts to protect cartilage in situations of trauma or challenge. These data can facilitate clarity around critical drivers, likely derived from adipose tissue, which may be foundational in the onset of OA with obesity or joint injury, as the basis for novel therapeutic strategies for OA.

## Conclusion

Adipose tissue is a critical antagonist of cartilage health and integrity independent of body mass, systemic inflammation, joint loading due to muscle weakness, and subchondral bone sclerosis. Specifically, these data implicate several context-relevant therapeutic targets: a protective role for inducing systemic increases in IL-10, TIMP-1, or systemically inhibiting IL-6/IL-12 cross-talk, leptin, IL-17, and CCL-17 to protect against cartilage knee joint damage. Characterizing adipose-cartilage crosstalk and signaling may lead to novel therapeutic opportunities for OA as well as other metabolic diseases involving adipose dysfunction. The fat-free LD mouse provides a powerful tool to disentangle the mechanisms involved in cartilage protection and vulnerability, and our ongoing efforts aim to clarify this mechanism by interrogating specific adipokines in LD mice, as well as the cytokines identified in the present study.

## Materials and Methods

### Animals and Design

All experimental procedures were approved by the Institutional Animal Care and Use Committee and were carried out in accordance with the guidelines prescribed by the Washington University School of Medicine Department of Comparative Medicine. LD mice and WT littermate controls were developed by crossing adiponectin-Cre mice with homozygous lox-stop-lox-ROSA-diphtheria toxin A mice (1) and were maintained at thermoneutrality throughout their life span (30C). Male and female offspring were weaned at 4-weeks of age to either a chow (10% kcal fat, standard chow) or high-fat (60% kcal fat, Research Diets, D12492) diet *ad libitum* through 28-weeks of age (n=7-16/group). Mice were group housed such that n=3-5/cage, and genotypes were mixed within a given cage. The timeline for these studies is depicted in *Supplementary Figure 1*. In the transplantation studies, male and female LD mice received either a mouse embryonic fibroblast transplant (MEF-rescue, (*17*) between 3-5 weeks of age, or wildtype fat transplant containing visceral and subcutaneous fat (WT fat-rescue (*14, 15*)) between 6-8 weeks of age. At 16-weeks, surgical animals underwent surgery for destabilization of the medial meniscus (DMM) in their left knee joints, and the right limb served as a non-surgical contralateral control (*6, 9, 10, 41*). Behavioral and metabolic assays were conducted throughout the course of this study. Mice were sacrificed at 28 weeks of age to assess OA severity, bone microstructure, and serum and synovial fluid inflammatory profiles.

### Body Weight and Composition

Mice were weighed weekly throughout the course of the study. Body fat was measured by dual-energy x-ray absorptiometry (DXA, Lunar Piximus) at 15 weeks and 27 weeks of age. In a subset of animals, an EchoMRI 3-in-1 instrument (Echo Medical Systems) was used to quantify body fat.

### Fat Transplantation

Mouse embryonic fibroblasts were harvested from E14 C57BL/6J embryos, resuspended in PBS, and delivered as an injection subcutaneously on the sternal aspect of the animals, as described previously (*17*). Embryos were isolated immediately after the pregnant dams were cervically dislocated. The liver, heart and intestine were removed from decapitated embryos, and the remaining tissue was minced and digested in 1.25mL 0.05% trypsin for 45min in a 37°C water bath. Trypsin was neutralized with a mixture of DMEM and 10% fetal bovine serum, triturated vigorously by pipetting up and down ~30 times, and filtered using a 70μm cell strainer. The filtrate was then centrifuged at 500g for 6 minutes and the cell pellet was resuspended in 250μL PBS. The injection was delivered using a 27-gauge insulin syringe into a donor LD mouse that was 3-5 weeks old, and anesthetized under 2-3% isoflurane.

Mature fat depots were implanted in accordance with previously described methods (*14, 15*). Donor WT mice (6-8 weeks old) were euthanized and subcutaneous and visceral fat pads were harvested and cut into 100-150 mg pieces, in PBS on ice. Recipient mice were anaesthetized using 2-3% isoflurane, dorsal hair was shaved, fat was implanted (~1g/mouse), and incisions were closed with absorbable sutures.

### Post-Traumatic OA Induction by Destabilization of the Medial Meniscus

At 16-weeks of age, mice were challenged with unilateral destabilization of the medial meniscus on their left knee joint, in accordance to previously described methods (*10, 41*). Briefly, anaesthetized mice were positioned supine in a custom-designed device that maintained 90 degrees of knee flexion in the left knee. The medial meniscotibial ligament was approached from the medial aspect of the knee by creating a small incision in the skin, followed by an incision in the joint capsule, and ultimately transected. The joint capsule and skin were closed by absorbable sutures and Vetbond (3M, Inc.) skin glue.

### Histological Assessment of OA, Synovitis, and Osteophytes

At 28-weeks of age, 12-weeks post DMM surgery, mice were sacrificed and knee joints were assessed using histology. Knee joints were harvested, all excess tissue and muscle were trimmed, and fixed in 4% paraformaldehyde for 24hrs. Limbs were decalcified in 10% Formic Acid (Cal-Ex) for 72hrs, dehydrated, and processed, and embedded in paraffin blocks. Serial frontal plane sections were cut at 5μm and stained with either hematoxylin, fast green, and safranin O (saf-O/fast green), or hematoxylin and eosin (H&E). Saf-O/Fast Green slides were scored using Modified Mankin Criteria to operationalize knee joint damage (*6, 10*). This scoring system employs a 30-point system to four compartments of the frontal plane section: medial tibial plateau, lateral tibial plateau, medial femoral condyle, lateral femoral condyle which is added to generate the total Modified Mankin Score for a joint that serves as the semiquantitative measure for joint damage. To assess synovitis in these joints, H&E stained slides were graded for synovial stroma density (0-3) and synovial lining thickness (0-3) for all four compartments of the synovium in the joint, corresponding to the four regions assessed in the Mankin Score. These values were summed to provide a total semiquantitative measure of synovitis (*6, 10*). Finally, to determine whole joint osteophytosis, the same four quadrants of each knee joint were graded from 0-3 using previously described methods (*10*), and summed for a cumulative score per joint. All histological scores represent the average values from scores of three blinded graders, and the interrater reliability measured by intraclass correlation was r>0.90, p<0.001.

### Bone Microstructural Analysis

Whole knee joints were scanned by micro-computed tomography (Bruker SkyScan1176) at a resolution of 18μm isotrophic voxel resolution according to previously reported methods (*6, 9*). To reduce beam hardening, a 0.5mm aluminum filter was used during scanning. Hydroxyapatite calibration phantoms were scanned to calibrate bone density. Scans were reconstructed to 3D images using NRecon software, and CTAn Software was used to segment subchondral and trabecular regions from the medial tibial plateau, lateral tibial plateau, medial femoral condyle, and lateral femoral condyle for analysis. The tibial epiphysis was identified using the subchondral plate and growth plate as references. The tibial metaphysis was defined as the 1-mm area directly below the growth plate. The main outcomes reported from microCT images are bone mineral density (BMD), bone volume/total volume BV/TV (%), trabecular number, and trabecular thickness.

### Serum and Synovial Fluid Profiling

All mice were fasted prior to sacrifice. Serum was collected in a serum separator tube and allowed to clot at room temperature. Synovial fluid was collected using an alginate pad that was digested in alginate lyase until a stop solution was applied (*6, 9*). Both fluids were stored at −80°C until analysis by Luminex multiplex 44-plex chemokine/cytokine array assay (Eve Technologies, Calgary, AB). The 44-plex assay included: Eotaxin, Erythropoietin, 6Ckine, Fractalkine, G-CSF, GM-CSF, IFNB1, IFNγ, IL-1α, IL-1β, IL-2, IL-3, IL-4, IL-5, IL-6, IL-7, IL-9, IL-10, IL-11, IL-12 (p40), IL-12 (p70), IL-13, IL-15, IL-16, IL-17, IL-20, IP-10, KC, LIF, LIX, MCP-1, MCP-5, M-CSF, MDC, MIG, MIP-1α, MIP-1β, MIP-2, MIP-3α, MIP-3B, RANTES, TARC, TIMP-1, TNFα, and VEGF. Serum was assessed at a concentration of 1:2, synovial fluid was evaluated at a concentration of 1:75 based on the minimum concentration of 75 μL.

ELISA for serum leptin was conducted using a mouse/rat Leptin Quantikine ELISA Assay kit (R&D Systems, MOB00) at a concentration of 1:2 in LD mice, and all other groups were evaluated at the manufacturer’s recommended 1:20 dilution. The lower limit of detection of the assay was 22 pg/mL, and the Inter-Assay variability was 3-4%.

### Insulin and Glucose Tolerance Tests

Insulin and glucose tolerance tests were performed 4 weeks post-DMM surgery (20 weeks of age) after fasting mice for a minimum of 4 hours. Each test was conducted at least one week apart in all animals. For both tests, fasting glucose levels were measured by tail bleed at timepoint zero to establish a baseline. For glucose tolerance tests, animals were administered 1 mg/kg (10% dextrose: 1% volume/body mass) by IP injection, and for insulin tolerance tests, 0.75 U/kg body mass of insulin (Humulin R diluted to 75mU/ml, 1% volume/body mass) was administered by IP injection (*17*). Serial blood glucose measurements were taken via tail vein at 20, 40, 60, and 120 minutes after injection with a glucose meter (Contour®, Bayer).

### Metabolic Parameters

Indirect calorimetry and activity were measured over 36 hours after a 6-hour acclimatization period using a Phenomaster (TSE Systems) Metabolic Caging System at room temperature 22°C at 22-24 weeks of age. Each mouse was individually housed during the observation period and given free access to food and water. Data were analyzed using CalR web-based tool for indirect calorimetry experiments (*42*) using the two group or four group templates, and 1-way ANOVA with Tukey’s post-hoc test.

### Pain and Behavioral Testing

All animals were acclimatized to all equipment 1 day prior to the onset of testing. Forelimb grip strength was used to measure muscle strength both prior to DMM at 15-weeks of age, and prior to sacrifice at 27-weeks of age, using a Chatillon DFE Digital Force Gauge (Johnson Scale Co.). Each mouse was evaluated 5 times with a minimal resting period of 90s between trials, and these trials were averaged to determine the grip strength value for each mouse. Two measures for pain were conducted at 27-weeks of age. To assess tactile allodynia in the DMM limb, an Electronic Von Frey assay was used. Mice were placed individually in acrylic cages (12 × 10 × 17 cm high) with a wire grid floor. Hind paws of DMM limbs were stimulated 2-3 times. A hand-held force transducer with a 0.5 mm 2 polypropylene tip (Electronic Von Frey anesthesiometer, IITC Inc., Life Science Instruments, Woodland Hills, CA, USA) was used to apply force against the central aspect of the DMM hind paw. The intensity of the stimulus (g) was recorded by the tester when the paw was withdrawn. The stimulation of the paw was repeated until the animal presented three similar measurements. To assess mechanical hyperalgesia, pressure-pain tests were conducted using a Small Animal Algometer (SMALGO, Bioseb, Pinellas Park, FL, USA) (*6*). Three trials of the surgical and nonsurgical limb were collected by applying a steadily increasing force to the lateral aspect of each limb until the limb was withdrawn. The average of three trials for each limb was reported, and maximum value of 450g was employed to avoid tissue damage to the knee joint.

### Statistical Analysis

Statistical analysis and number of animals/group for each experiment is described in each figure caption. Analysis was performed in Graphpad Prism 8 (Graphpad Software, La Jolla, Ca) or SPSS Statistics 23 (IBM). *A priori* (was defined as 0.05. Data were presented as mean±standard error, and were compared by either 1-way, 2-way (group*limb, genotype*sex, genotype*diet) or 3-way (limb*genotype*diet; limb*genotype*sex) ANOVA with Dunnet’s, Sidak’s or Tukey’s post-hoc test.

## Supplementary Materials

## General

The authors with to thank Natalie Kelly and Erica Ely for assistance with behavior testing experiments, and Sara Oswald for her assistance with editing and figure composition.

## Author contributions

KHC and FG conceived the studies. KHC, KLL, YRC, ST, GAM, CAH, FG, CTNP assisted with surgical procedures and adipose tissue implants. KLL, ENP, DF, IH, LES, AKO, RHT, CTNP assisted with animal experiments and data analysis. All authors contributed to the integration and interpretation of the data. KHC and FG wrote the manuscript. All authors approved the draft and provided critical feedback to final version.

## Data and materials availability

All data needed to evaluate the conclusions in the paper are present in the paper and/or the Supplementary Materials. Additional data related to this paper may be requested from the authors under an MTA.

---

**Supplementary Figure 1.**
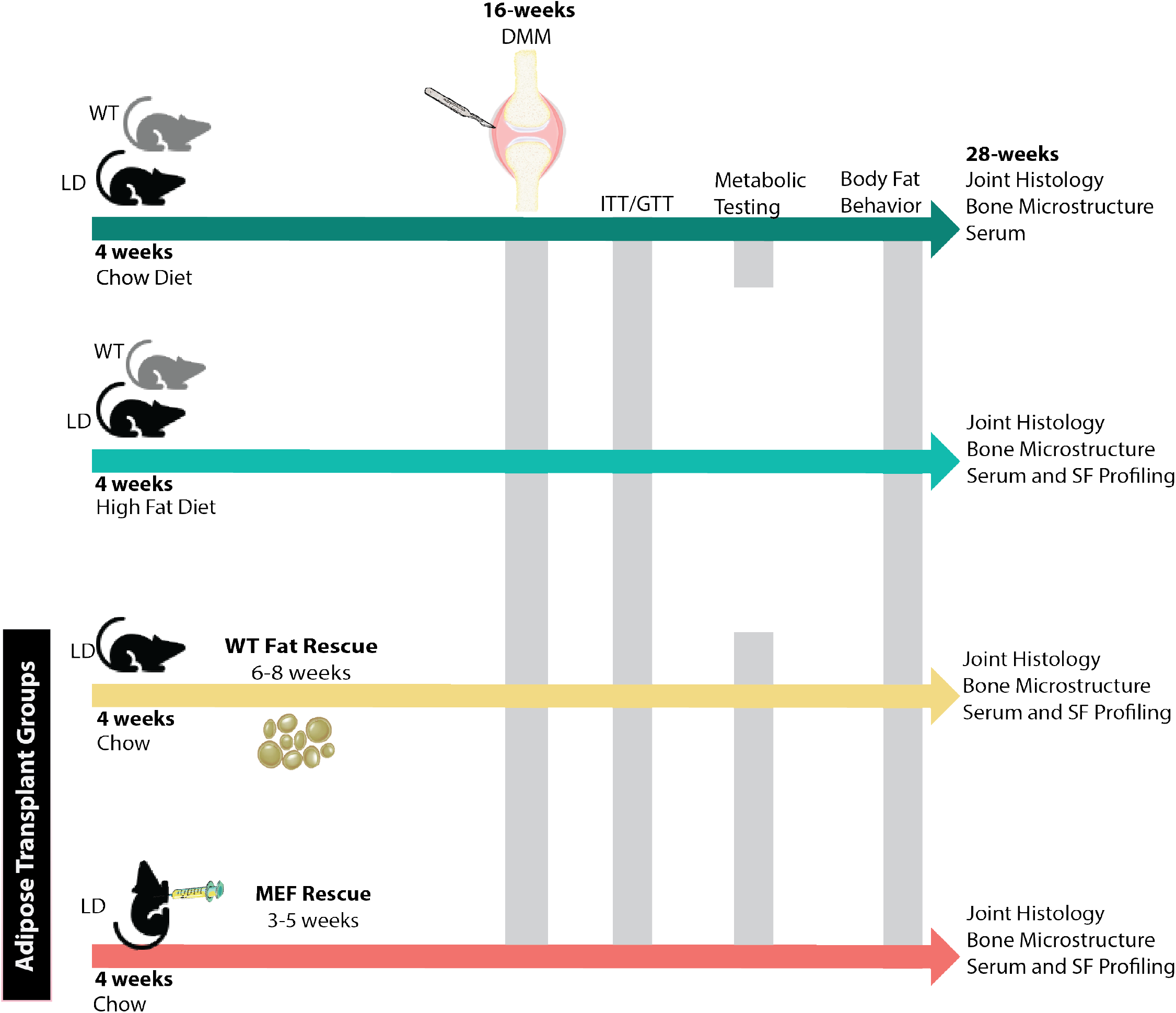
Timeline for study. First, male and female LD mice and WT littermate controls were fed a chow-control diet from weaning (4 weeks of age) and were challenged with DMM surgery at 16 weeks of age. Metabolic profiles were determined using insulin tolerance tests (ITT) and glucose tolerance tests (GTT) at 20 weeks of age, 4-weeks post DMM. At 22-24 weeks of age, metabolic testing using indirect calorimetry assessments were completed using a Phenomaster device. At 27-weeks of age, one week prior to sacrifice, body composition and behavior assays were completed. Animals were euthanized at 28-weeks of age, 12-weeks post DMM surgery. Tissues were preserved for joint histology and bone microstructure, and serum, synovial fluid, and fecal matter was stored at −80C for profiling. The same timeline was followed for male and female LD and WT HFD-fed mice, with the exception of indirect calorimetry metabolic testing – these animals were not evaluated using the Phenomaster. In the transplant groups, LD host mice either received wild type fat rescue at 6-8 weeks of age, or a cell-based injection of preadipocyte mouse embryonic fibroblasts at 3-5 weeks of age. Both transplant groups were maintained on chow diets, and followed the same protocol as the chow-fed LD and WT mice. The same euthanization and tissue preparation procedures were employed to all groups of mice at 28-weeks of age.

**Supplementary Figure 2.**
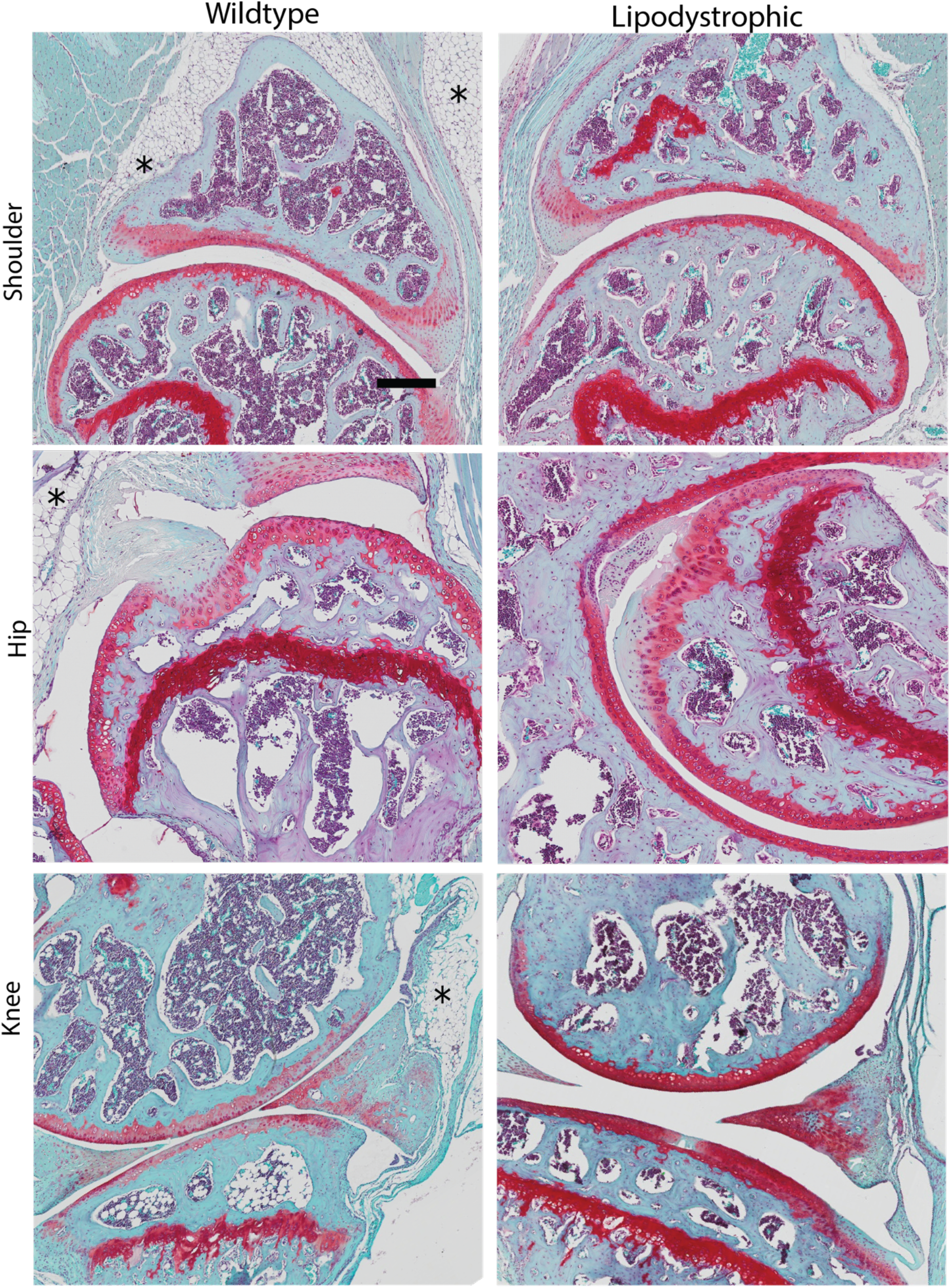
Fat pads are absent in Shoulder, Hip, and Knee Joints of LD mice. LD mice do not demonstrate fat pads adjacent to shoulder, hip, or knee joints, indicated with black asterisks in the left panels. Scale bar indicates 100μm.

**Supplementary Table 1.** Serum Changes in Chow and HFD Male Mice by Genotype, data are shown as Mean ± SEM.

|  | **Chow** |  | **HFD** |  | **Main Effects (p<0.05)** |
| --- | --- | --- | --- | --- | --- |
| ***Pro-Inflammatory Mediators*** | WT | LD | WT | LD |  |
| IL-1α (pg/mL) | 140.7±20.9 | 368.7±31.6<sup>a</sup> | 294.0±28.2<sup>a</sup> | 243.38±39.5<sup>a</sup> | Genotype, Diet*Genotype |
| IL-1β (pg/mL) | 5.4±1.1 | 38.3±13.9<sup>a</sup> | 8.3±3.5 | 23.4±8.0<sup>a</sup> | Genotype |
| IL-6 (pg/mL) | 3.8±1.3 | 14.3±4.2<sup>a</sup> | 2.7±0.9 | 11.4±2.65<sup>a</sup> | Genotype |
| TNF- $\alpha$ (pg/mL) | 9.1±1.8 | 25.6±5.7<sup>a</sup> | 6.5±2.2 | 17.8±3.7<sup>a</sup> | Genotype |
| IP-10 / CXCL-10 (pg/mL) | 114.4±11.5 | 155.2±20.0<sup>a</sup> | 127.1±10.0 | 157.5±11.1<sup>a</sup> | Genotype |
| MCP-1 (pg/mL) | 64.5±9.4 | 141.9±27.2<sup>a</sup> | 89.7±20.3 | 117.5±35.5<sup>a</sup> | Genotype |
| IL-12p70 (pg/mL) | 83.2±22.9 | 310.4±136.2<sup>a</sup> | 80.9±22.7 | 250.8±77.3<sup>a</sup> | Genotype |
| IL-15(pg/mL) | 70.6±6.0 | 75.3±14.6 | 154.9±31.0 | 1140.9±415.8<sup>ab</sup> | Diet, Genotype, Diet*Genotype |
| IL-17 (pg/mL) | 5.8±1.1 | 7.6±1.6 | 6.7±1.2 | 14.0±4.4 | NS |
| IFN β-1 (pg/mL) | 157.2±10.4 | 166.9±30.9 | 244.5±36<sup>a</sup> | 212.8±32.3<sup>a</sup> | Diet |
| TARC / CCL-17 (pg/mL) | 19.0±3.0 | 35.0±6.3<sup>a</sup> | 21.7±2.4 | 18.3±2.3 | Diet p=0.09, Diet*genotype |
| ***Anti-Inflammatory Mediators*** |  |  |  |  |  |
| IL-10 (pg/mL) | 7.2±1.8 | 81.0±27.9<sup>a</sup> | 11.4±4.0 | 74.4±16.2<sup>a</sup> | Genotype |
| IL-4 (pg/mL) | 1.3±0.3 | 5.7±1.5<sup>a</sup> | 1.7±0.4 | 3.7±1.5<sup>a</sup> | Genotype |
| TIMP-1(ng/mL) | 0.8±0.1 | 2.1±0.2<sup>a</sup> | 1.6±0.<sup>a</sup> | 3.7±0.3<sup>ab</sup> | Diet, Genotype |

Sidak's Post-hoc Test
<sup>a</sup>p<0.05 compared to WT Chow
<sup>b</sup>p<0.05 compared to WT HFD
n=10-14/group

**Supplementary Table 2.** MicroCT outcomes for all groups: Bone Mineral Density (BMD), Bone Volume/Total Volume (BV/TV), Trabecular Number, and Trabecular Thickness. Data were compared in 3 groups: Chow Male and Female LD and WT) by 3-way ANOVA (limb, sex, genotype), HFD Male and Chow Male LD and WT (limb, diet, genotype), and MEF rescue (MEF-R) and WT fat rescue (WF-R) vs. Chow LD and WT, 2-Way ANOVA (limb, group). Data are separated into contralateral non-surgical limbs (2a) and DMM surgical limbs (2b) for viewing purposes.

**2a. Contralateral Non-Surgical Limbs**
|  |  | Chow |  |  |  | Main Effects | HFD |  | Main Effects | Transplant |  | Main Effects |
| --- | --- | --- | --- | --- | --- | --- | --- | --- | --- | --- | --- | --- |
|  |  | Female |  | Male |  | M V. F<br>P<0.05 |  |  | HFD M V. Chow M<br>P<0.05 |  |  | Transplant V. Chow<br>P<0.05 |
| Outcome | Region | WT | LD | WT | LD |  | WT | LD |  | WF-R | MEF-R |  |
| *BMD*<br>(gCaHA/cm<sup>3</sup>) | LTP | 0.62±0.02 | 0.84±0.02<sup>ab*</sup> | 0.66±0.02 | 0.72±0.02 | Sex, Genotype,<br>Sex*Genotype | 0.66±0.02 | 0.71±0.04 | Genotype | 0.75±0.03 | 0.71±0.4 | Group |
|  | LFC | 0.52±0.01 | 0.74±0.04<sup>a</sup> | 0.58±0.01 | 0.68±0.04 | Genotype | 0.58±0.13 | 0.64±0.02 | Genotype | 0.72±0.04<sup>a</sup> | 0.70±0.04 | Group |
|  | MFC | 0.51±0.02 | 0.75±0.05<sup>a</sup> | 0.63±0.02 | 0.71±0.04 | Genotype,<br>Sex*Genotype | 0.67±0.02 | 0.70±0.03 | Genotype | 0.77±0.03<sup>a</sup> | 0.75±0.05 | Group |
| *BV/TV (%)* | LTP | 49.2±1.0 | 85.8±2.1<sup>a*</sup> | 61.6±1.9 | 70.0±2.7 | Genotype,<br>Sex*Genotype,<br>Limb*Sex*Genotype | 56.1±2.1 | 67.1±2.6<sup>c</sup> | Diet, Genotype | 69.4±4.0 | 70.5±5.2 | Group |
|  | LFC | 41.5±0.3 | 73.0±4.3<sup>a</sup> | 52.9±1.4<sup>*</sup> | 58.4±2.3 | Genotype,<br>Sex*Genotype,<br>Limb*Sex*Genotype | 48.7±1.3 | 60.0±1.8<sup>c</sup> | Genotype | 64.4±4.4 | 64.9±4.6 | Group |
|  | MFC | 42.4±0.4 | 77.3±4.5<sup>a</sup> | 58.2±2.5 | 63.0±2.5<sup>a</sup> | Genotype,<br>Sex*Genotype | 58.9±1.6 | 66.1±2.1 | Genotype,<br>Limb*Genotype | 68.2±3.2 | 70.8±4.8<sup>a</sup> | Group |
| *Trabecular Thickness*<br>(mm) | MTP | 0.08±0.01 | 0.08±0.01 | 0.09±0.01 | 0.08±0.01 | None | 0.09±0.01 | 0.08±0.01 | None | 0.09±0.01 | 0.07±0.01 | None |
|  | LTP | 0.09±0.001 | 0.09±0.007 | 0.09±0.005 | 0.09±0.006 | Sex*Genotype | 0.08±0.005 | 0.09±0.004 | Diet, Genotype | 0.11±0.004 | 0.09±0.009 | Group |
|  | LFC | 0.10±0.01 | 0.10±0.01 | 0.10±0.01 | 0.10±0.01 | Genotype, Sex*Limb | 0.10±0.01 | 0.10±0.01 | Genotype,<br>Limb*Diet | 0.10±0.01 | 0.10±0.01 | Limb*<br>Group |
|  | MFC | 0.10±0.01 | 0.11±0.01 | 0.10±0.01 | 0.11±0.01 | Genotype | 0.11±0.01 | 0.11±0.01 | Limb, Genotype,<br>Limb*Diet,<br>Limb*Genotype | 0.12±0.01 | 0.11±0.01 | Limb |
|  | TM | 0.05±0.01 | 0.07±0.01<sup>*</sup> | 0.06±0.01 | 0.05±0.01 | Limb, Genotype,<br>Limb*Sex,<br>Limb*Sex*Genotype | 0.06±0.01 | 0.06±0.01 | Limb, Limb*Diet,<br>Limb*Genotype,<br>Limb*Diet*<br>Genotype | 0.07±0.01<sup>b</sup> | 0.07±0.01<sup>b</sup> | Limb*<br>Group |
| *Trabecular Number*<br>(1/mm) | MTP | 6.9±0.3 | 11.6±1.3<sup>a</sup> | 7.4±0.4 | 8.9±0.5 | Genotype,<br>Sex*Genotype | 7.3±0.2 | 9.7±1.0 | Diet | 10.0±1.0 | 11.5±1.3 | Group |
|  | LTP | 6.6±0.6 | 9.1±0.7 | 6.5±0.2 | 8.0±0.7 | Limb, Genotype | 7.6±0.6 | 6.8±0.2 | Genotype | 6.5±0.2 | 8.6±0.8<sup>Ad</sup> | Group |
|  | LFC | 4.4±0.1 | 6.8±0.4<sup>a*</sup> | 5.6±0.2<sup>*</sup> | 5.7±0.2 | Genotype,<br>Sex*Genotype,<br>Limb*Sex*Genotype | 5.1±0.1 | 5.7±0.1<sup>c</sup> | Genotype,<br>Limb*Diet*<br>Genotype | 6.2±0.5 | 5.9±0.3 | Limb*<br>Group |
|  | MFC | 4.6±0.2 | 6.8±0.4<sup>a*</sup> | 5.5±0.2 | 5.9±0.1 | Genotype<br>Limb*Genotype<br>Sex*Genotype<br>Limb*Sex*Genotype | 5.6±0.1 | 5.8±0.1 | Genotype | 5.6±0.2 | 6.7±1.0<sup>a</sup> | Group P=0.07 |
|  | TM | 1.2±0.3 | 3.5±0.8<sup>a*</sup> | 0.8±0.1 | 1.8±0.3 | Genotype, Limb*Sex,<br>Limb*Sex*Genotype | 1.0±0.2 | 1.8±0.3 | Limb, Genotype,<br>Limb*Diet,<br>Limb*Genotype | 2.0±0.2 | 2.2±0.4 | Limb, Group |

**2b. DMM Surgical Limbs MicroCT Outcomes.**
|  |  | Chow |  |  |  | Main Effects | High-Fat Diet Males |  | Main Effects | Fat Transplant |  | Main Effects (P<0.05) |
| --- | --- | --- | --- | --- | --- | --- | --- | --- | --- | --- | --- | --- |
|  |  | Female |  | Male |  | M V. F<br>P<0.05 | Genotype |  | HFD M V. Chow M<br>P<0.05 |  |  | Transplant V.<br>Chow M |
| Outcome | Region | WT | LD | WT | LD |  | WT | LD |  | WF-R | MEF-R |  |
| *BMD*<br>(gCaHA/cm<sup>3</sup>) | LTP | 0.61±0.03 | 0.90±0.02<sup>ab*</sup> | 0.64±0.03 | 0.74±0.04 | Sex, Genotype,<br>Sex*Genotype | 0.63±0.02 | 0.71±0.03 | Genotype | 0.75±0.04 | 0.70±0.04 | Group |
|  | LFC | 0.52±0.01 | 0.70±0.06<sup>a</sup> | 0.56±0.02 | 0.67±0.04 | Genotype | 0.56±0.01 | 0.66±0.02<sup>c</sup> | Genotype | 0.64±0.04 | 0.69±0.05 | Group |
|  | MFC | 0.56±0.03 | 0.79±0.08<sup>a</sup> | 0.61±0.03 | 0.75±0.06 | Genotype, Sex*Genotype | 0.64±0.02 | 0.77±0.03 | Genotype | 0.70±0.02 | 0.77±0.06<sup>a</sup> | Group |
| *BV/TV (%)* | LTP | 53.5±2.9<sup>b</sup> | 74.8±11.1<sup>a</sup> | 61.3±2.5<sup>b</sup> | 73.5±1.8<sup>a</sup> | Genotype, Sex*Genotype,<br>Limb*Sex*Genotype | 55.0±2.1 | 66.1±2.1<sup>c</sup> | Diet, Genotype | 70.0±4.9 | 69.1±4.4 | Group |
|  | LFC | 43.0±0.1 | 66.5±8.6<sup>a</sup> | 52.5±1.8 | 62.5±2.4 | Genotype, Sex*Genotype,<br>Limb*Sex*Genotype | 48.7±1.0 | 59.7±2.2<sup>c</sup> | Genotype | 55.2±5.0 | 64.0±4.9 | Group |
|  | MFC | 43.7±1.7 | 71.4±9.4<sup>a</sup> | 56.9±1.7<sup>*</sup> | 71.0±3.0<sup>a</sup> | Genotype, Sex*Genotype | 55.2±1.5 | 70.1±2.2<sup>c</sup> | Genotype, Limb*Genotype | 61.8±2.6 | 73.1±5.3<sup>a</sup> | Group |
| *Trabecular Thickness (mm)* | MTP | 0.09±0.01 | 0.08±0.01 | 0.08±0.01 | 0.08±0.01 | None | 0.08±0.01 | 0.07±0.06 | None | 0.08±0.01 | 0.07±0.01 | None |
|  | LTP | 0.10±0.003 | 0.08±0.001 | 0.09±0.002 | 0.10±0.003 | Sex*Genotype | 0.09±0.003 | 0.09±0.004 | Diet, Genotype | 0.09±0.001 | 0.08±0.01<sup>b</sup> | Group |
|  | LFC | 0.10±0.01 | 0.10±0.01 | 0.10±0.01 | 0.11±0.01<sup>^†</sup> | Genotype, Sex*Limb | 0.09±0.01 | 0.10±0.01<sup>c</sup> | Genotype, Limb*Diet | 0.10±0.01 | 0.09±0.01<sup>b</sup> | Limb*Group |
|  | MFC | 0.10±0.01 | 0.12±0.01 | 0.11±0.01 | 0.13±0.01<sup>†</sup> | Genotype | 0.11±0.01 | 0.12±0.01<sup>c</sup> | Limb, Genotype, Limb*Diet,<br>Limb*Genotype | 0.12±0.01 | 0.12±0.01 | Limb |
|  | TM | 0.06±0.01 | 0.08±0.01 | 0.06±0.01 | 0.07±0.01<sup>†</sup> | Limb, Genotype,<br>Limb*Sex,<br>Limb*Sex*Genotype | 0.06±0.01 | 0.06±0.01<sup>b</sup> | Limb, Limb*Diet,<br>Limb*Genotype,<br>Limb*Diet*Genotype | 0.07±0.01 | 0.06±0.01<sup>b</sup> | Limb*Group |
| *Trabecular Number (1/mm)* | MTP | 6.2±0.4 | 11.1±2.4<sup>a</sup> | 7.5±0.4 | 9.5±0.6 | Genotype, Sex*Genotype | 8.7±0.4 | 12.5±1.7 | Diet | 12.5±3.1 | 13.0±3.4 | Group |
|  | LTP | 5.3±0.1 | 7.4±0.4 | 6.7±0.4 | 6.9±0.4 | Limb, Genotype | 6.0±0.1 | 7.1±0.4 | Genotype | 6.8±0.4 | 8.2±0.7 | Group |
|  | LFC | 4.7±0.1 | 6.3±0.3<sup>a</sup> | 5.3±0.2 | 5.7±0.2 | Genotype, Sex*Genotype,<br>Limb*Sex*Genotype | 5.3±0.1 | 5.7±0.1 | Genotype, Limb*Diet*<br>Genotype | 5.3±0.4 | 6.6±0.4<sup>ad</sup> | Limb*Group |
|  | MFC | 4.9±0.2 | 6.1±0.3<sup>†</sup> | 5.3±0.2 | 6.0±0.1<sup>a</sup> | Genotype<br>Limb*Genotype<br>Sex*Genotype<br>Limb*Sex*Genotype | 5.2±0.1 | 5.7±0.1 | Genotype | 5.2±0.2 | 6.0±0.3 | Group P=0.07 |
|  | TM | 1.0±0.2 | 3.1±0.8 | 0.9±0.1 | 3.1±0.4<sup>^†</sup> | Genotype, Limb*Sex,<br>Limb*Sex*Genotype | 0.9±0.2 | 2.0±0.2<sup>cb</sup> | Limb, Genotype, Limb*Diet,<br>Limb*Genotype | 2.1±0.3 | 3.0±1.0<sup>a</sup> | Limb, Group |

Sidak's Post hoc for 3-way ANOVAS, Tukey's post-hoc for 2-way ANOVA
<sup>a</sup>p<0.05 compared to WT chow control,
<sup>b</sup>p<0.05 vs. LD-M chow,
<sup>c</sup>p<0.05 compared to WT HFD control,
<sup>d</sup>p<0.05 WT fat transplant
\*p<0.05 by sex,
<sup>†</sup>p<0.05 within group by limb,
male n= 9-17/group, female n=5-12
**Abbreviations:** LTP: lateral tibial plateau, LFC: lateral femoral condyle, MFC: Medial femoral condyle; MTP: Medial tibial plateau, TM: Tibial Metaphysis; BMD: Bone Mineral Density, BV/TV: Bone Volume/ Total Volume

**Supplementary Table 3.** Male Synovial Fluid Profiles from Chow and High Fat Diet-fed LD and WT Mice.

|  | Chow |  |  |  | HFD |  |  |  | Main Effects<br>(p<0.05) |
| --- | --- | --- | --- | --- | --- | --- | --- | --- | --- |
| Genotype | WT |  | LD |  | WT |  | LD |  |  |
| (pg/mL) | Con | DMM | Con | DMM | Con | DMM | Con | DMM |  |
| IL-7 | 67.2±13.0 | 30.2±9.7 | 50.0±12.3 | 39.7±9.0 | 57.2±19.5 | 28.7±8.7 | 37.2±9.4 | 30.4±8.7 | Limb |
| Fractalkin<br>e | 8.7±0.3 | 9.3±1.0 | 8.5±0.5 | 8.0±0.4 | 11.4±1.0 | 11.3±1.0 | 10.2±0.8 | 9.5±0.8 | Limb, Diet |
| G-CSF | 44.7±6.3 | 36.4±8.1 | 63.5±10.3 | 63.1±12.2 | 32.7±7.1 | 40.5±9.3 | 38.8±7.4 | 54.4±9.3 | Genotype |
| IL-6 | 287.4±102.<br>2 | 176.4±96.2 | 230.1±89.7 | 978.5±427.4 | 622.8±257.<br>9 | 441.4±155.3 | 1176.4±615.1 | 1859.6±770.0 | Genotype |
| IL-12p40 | 226.8±78.0 | 259.9±86.0 | 189.4±52.3 | 103.5±33.8 | 313.1±66.9 | 305.6±55.0 | 138.6±27.0 | 102.0±31.3 | Genotype |
| IL-15 | 489.1±78.6 | 472.6±59.1 | 357.9±78.0 | 405.2±70.3 | 549.5±58.7 | 415.3±64 | 311.7±43.4 | 408.6±53.6 | Genotype |
| IFN-γ | 87.3±24.7 | 105.8±28.8 | 77.7±17.2 | 91.2±19.9 | 77.5±14.5 | 75.3±23.2 | 23.7±9.3 | 31.7±12.9 | Diet, Genotype |
| IL-16 | 3.1±0.6 | 4.2±0.6 | **5.9±1.2<sup>a</sup>** | 4.4±0.6 | 1.8±0.5 | 2.2±0.5 | 3.0±0.3 | 3.3±0.5 | Diet, Genotype |
| MIP-2 | 9780±529 | 10059±584 | 9913±601 | 9962±334 | 10028±442 | 8468±615 | 7468±642 | 7720±436 | Diet, Genotype,<br>Diet*genotype |
| IL-1β | 146.7±28.6 | 115.4±28.5 | 85.1±20.0 | 117.3±25.9 | 140.5±9.0 | 73.5±19.3 | 85.1±19.1 | 92.0±23.0 | Limb*Genotype |
| MCP-5 | 101.3±22.3 | 110.0±4.4 | 150.8±19.5 | 105.2±12.0 | 48.7±7.2 | 147.0±35.1 | 97.4±13.0 | 105.9±20.6 | Limb*Genotype<br>, Diet*Genotype |
| MDC | 20.5±4.0 | 40.1±7.7 | **82.2±14.0<sup>a</sup>** | **75.7±23.0<sup>a</sup>** | 29.6±6.6 | **100.0±22.7<sup>a</sup>**<br>**b** | 21.4±4.8 | 27.8±4.7 | Limb,<br>Limb*Genotype<br>, Diet*Genotype |
| TNF-α | 9.4±1.0 | 14.0±2.3 | 11.0±2.2 | 9.3±1.5 | 6.6±0.7 | 6.7±1.3 | 5.6±1.0 | 5.7±1.1 | Diet |
| Eotaxin | 556.9±67.3 | 541.3±54.0 | 521.7±69.2 | 431.0±50.0 | 647.4±163.<br>4 | 639.0±94.0 | 679.9±115.0 | 860.4±236.0 | Diet |
| LIX | 618.3±169.<br>0 | 1268.9±292.<br>1 | 634.3±151.<br>0 | 1018.0±261.<br>0 | 520.7±164.<br>0 | 445.7±121.4 | 167.8±58.3 | 365.6±125.0 | Diet |
| IL-1α | 18.2±8.5 | 263.2±75.0 | 286.15±90.<br>3 | 109.1±30.0 | 881.7±163.<br>9 | 612.7±232.0 | 1135.9±347.0<br>a | 1475.7±478.6<br>a | Diet |
| IP-10 | 20.2±5.1 | 18.4±3.3 | 31.8±6.2 | 32.2±5.8 | **53.3±10.8<sup>a</sup>** | **70.0±17.0<sup>a</sup>** | **51.3±6.1<sup>a</sup>** | **70.3±11.2<sup>a</sup>** | Diet |

Dunnet's Post-hoc Test
<sup>a</sup>p<0.05 compared to WT chow contralateral
<sup>b</sup>p<0.05 compared to WT HFD contralateral
n= 10-12/group

**Supplementary Table 4.** Serum Changes in WT Fat rescue (WF-R) and MEF rescue (MEF-R) Corrected Mice by Genotype, data are shown as Fold Change Mean ± SEM.

|  | **Chow** |  | **Fat-Transplant** |  | **1-way ANOVA p-value** |
| --- | --- | --- | --- | --- | --- |
| (Fold Change) | WT | LD | WF-R | MEF-R |  |
| MCP-5 | 1.1±0.1 | 1.0±0.2 | 2.2±0.3<sup>a</sup> | 1.9±0.3<sup>a</sup> | **0.002** |
| IL-15 | 1.0±0.2 | 1.0±0.2 | 1.9±0.6 | 3.6±1.1<sup>a</sup> | **0.015** |
| IFN β-1 | 1.0±0.3 | 0.8±0.1 | 21.5±8.8<sup>a</sup> | 7.4±2.3 | **0.011** |
| TARC / CCL-17 | 1.0±0.2 | 1.9±0.4<sup>a</sup> | 1.9±0.2<sup>a</sup> | 1.3±0.1 | **0.0187** |
| IL-4 (pg/mL) | 1.0±0.3 | 2.5±1.0 | 0.6±0.1 | 0.6±0.1 | **0.0420** |
| MCP-1 | 1.1±0.2 | 1.7±0.4 | 0.9±0.2 | 0.9±0.1 | 0.094 |
| IL-12p70 | 1.0±0.1 | 1.8±0.5 | 1.1±0.2 | 1.1±0.2 | 0.094 |
| LIF | 0.9±0.3 | 1.6±0.2 | 2.0±0.8 | 2.1±0.7<sup>a</sup> | 0.167 |
| IL-1β | 1.0±0.2 | 3.0±1.2<sup>a</sup> | 0.5±0.2 | 0.5±0.2 | 0.187 |
| IP-10 / CXCL-10 | 1.0±0.1 | 1.8±0.5 | 1.0±0.3 | 1.1±0.2 | 0.181 |

Dunnet's Post-hoc Test
<sup>a</sup>p<0.05 compared to WT
<sup>b</sup>p<0.05 compared to
n=10-14/group

**Supplementary Table 5.**
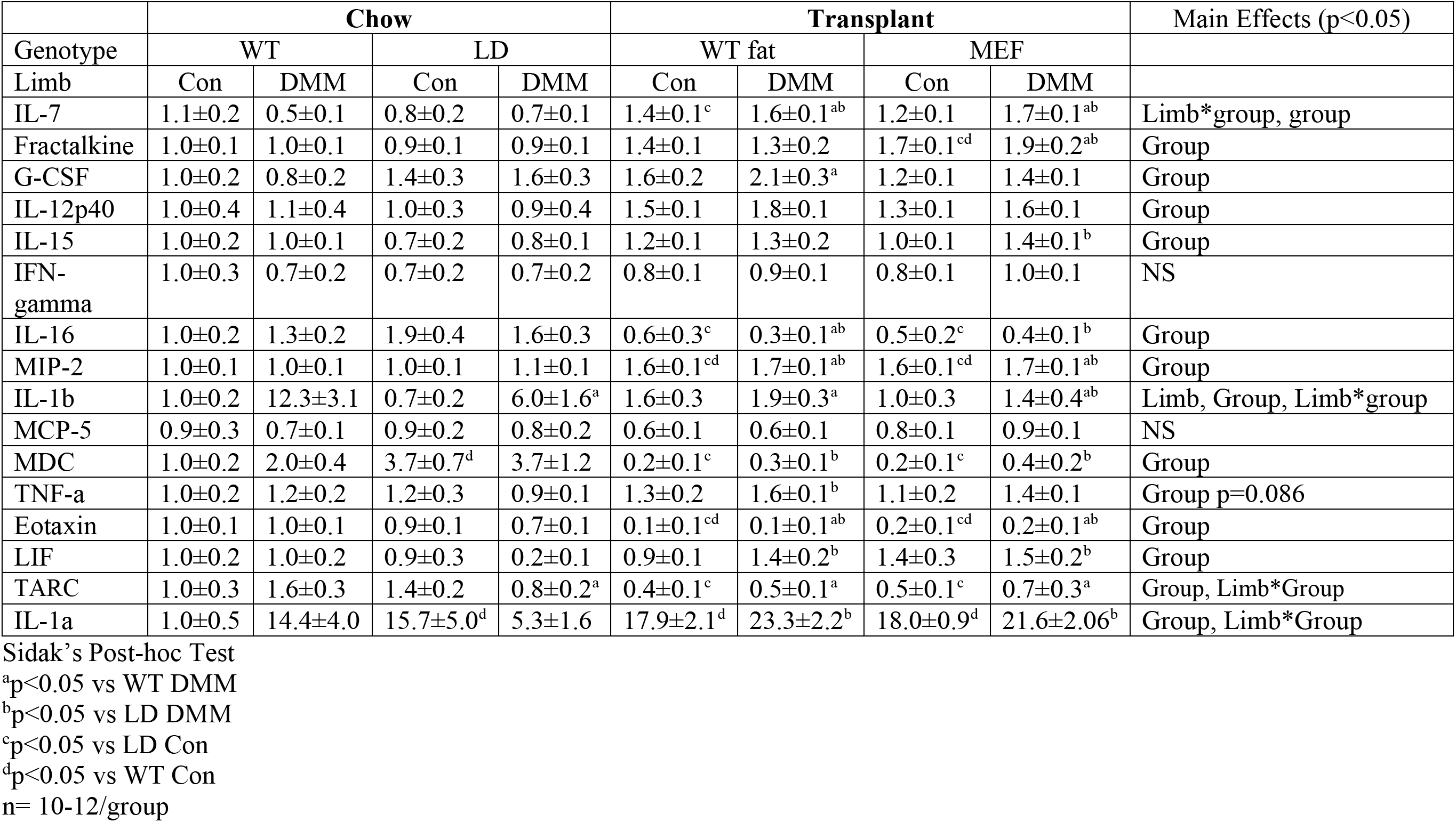
Synovial Fluid Changes in MEF/WT

